# Finding associations in a heterogeneous setting: Statistical test for aberration enrichment

**DOI:** 10.1101/2020.03.23.002972

**Authors:** Aziz M. Mezlini, Sudeshna Das, Anna Goldenberg

## Abstract

Most two-group statistical tests are implicitly looking for a broad pattern such as an overall shift in mean, median or variance between the two groups. Therefore, they operate best in settings where the effect of interest is uniformly affecting everyone in one group versus the other. In real-world applications, there are many scenarios where the effect of interest is heterogeneous. For example, a drug that works very well on only a proportion of patients and is equivalent to a placebo on the remaining patients, or a disease associated gene expression dysregulation that only occurs in a proportion of cases whereas the remaining cases have expression levels indistinguishable from the controls for the considered gene. In these examples with heterogeneous effect, we believe that using classical two-group statistical tests may not be the most powerful way to detect the signal. In this paper, we developed a statistical test targeting heterogeneous effects and demonstrated its power in a controlled simulation setting compared to existing methods. We focused on the problem of finding meaningful associations in complex genetic diseases using omics data such as gene expression, miRNA expression, and DNA methylation. In simulated and real data, we showed that our test is complementary to the traditionally used statistical tests and is able to detect disease-relevant genes with heterogeneous effects which would not be detectable with previous approaches.

## 1 Introduction

Two-group statistical tests are widely used to characterize significant differences associated with an intervention or a condition. In a case/control setting, these tests can pinpoint variables of interest in the dataset analysed. For example, gene expression data has been extensively used to characterize genes and pathways relevant to genetic diseases. If a gene is found to be differentially expressed (over-expressed or under-expressed) in the disease cases when compared to healthy controls then it can potentially be associated with the disease. The differentially expressed gene can be causal for the disease, in which case it can become a candidate for therapeutic intervention, or the association it can be non-causal: for example a compensatory or a downstream consequence of the disease state itself (immune reaction, treatment effect, etc.). Nevertheless, finding the differentially expressed genes often generates candidates that are further tested for their mechanistic involvement in the disease [7,40,77]. The typical approach for finding differentially expressed genes relies on statistical tests (e.g. Limma [52]) that look for a broad pattern such as a global shift in mean expression between a target group (the cases) and a control group.

In this paper, we look for another mode of association that does not present as the typical broad pattern of mean difference typically targeted by the widely used statistical tests. In this mode, the considered variable will contain a significant number of outliers in cases (compared to controls), while the remaining majority of the cases will not be distinguishable from controls. To use the gene expression example again, consider the scenario where 10% of disease cases have an extremely low level of expression for a gene of interest but only 1% of the controls do. In the rest of the paper, we will call this hypothesized pattern of association “aberration enrichment” to distinguish it from the broad pattern of a mean/median/variance difference between two groups targeted by the currently used approaches. We will also describe the features/genes exhibiting this aberration enrichment pattern as “aberration enrichment features” or “features with heterogeneous effects”.

There are many reasons to believe this mode of aberration enrichment exists and is particularly relevant for the characterization of complex diseases. **First**, in complex diseases, it is expected that the disease causes would be spread across multiple genes, such that any particular gene would only be causal in a small proportion of the patients. This remains likely even when multiple different causal genes need to be hit to reach the disease state (as it is known to be the case in cancer [68]). It is unlikely in a complex disease to observe a single <u>causal</u> gene or factor that can broadly separate cases and controls. If we observe a single factor where the value for most patients differs from the typical value in healthy controls then that factor is more likely to be a downstream consequence of the disease than to be causal. Otherwise, the disease would be mostly explained/caused by that one factor/gene contradicting the definition of a complex disease.

**Second**, work by major consortia have recently highlighted the importance of looking at rare events and outliers, rather than broad differences, to characterize disease biology. For example, work in the GTEX consortium [32] established links between being an outlier for a gene’s expression and having large impact rare cis-regulatory variants nearby. They also further linked the expression aberrations with diseases by selecting disease-associated variants and showing that they were highly enriched in variants predicted to generate expression outliers [32]. The pattern of aberration enrichment we are targeting in this paper corresponds to the expression outliers and could result from rare regulatory events involving SNVs, indels, structural and epigenetic variants such as the ones investigated in the GTEX paper [32].

**Third**, there are many known genetic diseases where a portion of patients is explained by aberrant or outlier levels of a variable of interest such as a gene’s expression [13,19,28,31]. In the examples cited, a gene is associated with the disease through the presence of harmful coding variants in a proportion of patients. The authors observed that there were more patients without coding variants but whose expression levels for that gene are abnormally low. These expression aberrations were observed by manually counting the number of individuals with extremely low expression in a suspected causal gene. Providing a statistical test for automatically detecting aberration enrichment can further empower and formalize such analyses. Although these observations were mainly done in rare or Mendelian diseases where the causal gene is known through proven causal coding variation, the same mode of action (presence of expression aberrations/outliers) could also be relevant in more complex diseases.

The reasoning we made here for gene expression data also holds for other types of quantitative omics data where an enrichment in aberrations is a possible relevant pattern for disease association. This includes miRNA and noncoding RNA expression, protein expression and DNA methylation.

In this paper, we present a new statistical test that aims at detecting novel associations through aberration enrichment: The presence of outlier values in a small but significant proportion of the cases. Outliers may be present in the data for many reasons including biological variation and technical artifacts and are not necessarily associated with a phenotype of interest. The focus here is not on outlier detection per se (such as in [6]) but on finding consistent aberrations (in the same direction) that are significantly enriched in a subset of cases when compared to controls. Genes/factors discovered through this pattern of aberration enrichment can shed light on novel mechanisms and disease subtypes [55] undetected by previous methods looking for broad signals.

This pattern of aberration enrichment is discussed in the literature under other names. For example, OSACC [51] aims to identify signals that are present in a subset of the cases. They look for the best subset of individuals that leads to a stronger SNP association compared to taking all cases and all controls. In their case, the subset selection is guided and defined by a continuous known covariate variable, such as age (the context is finding G*E associations). In clinical trials literature, the pattern is known as heterogeneous treatment effects [27] where a drug could be working well in a subset of patients but still fail to show efficacy when considering all participants because the statistical methods used are looking for a mean effect. This has previously been discussed as “the trouble with the averages” [30].

Using simulations, we show that our test is well calibrated and more powerful in detecting the aberration enrichment pattern compared to the widely used statistical tests, including t-test, Limma, Wilcoxon, Levene and Kolmogorov-Smirnov test. We then use our test to examine 12 real datasets from GEO [5] spanning various cancers, neurodegenerative and auto-immune conditions and 3 different data types (gene expression, miRNA expression and DNA methylation). We discover new meaningful disease associations that were not captured by the traditional approaches.

## 2 Results

### 2.1 Overview of our statistical test

The test presented in this paper is motivated by GSEA (Gene Set Enrichment Analysis) [43,64]. GSEA takes a ranked list of genes and an annotated gene set (for example a pathway) and tests if the set is enriched at the top or bottom of the ranked list. An enrichment score is iteratively computed while walking through the ranked list. The score is incremented every time a positive gene (from the set) is encountered and is decremented every time a negative gene is encountered. The maximum enrichment score is saved and its significance is assessed by a permutation test.

In our test, we compute a ranked list of samples (cases and controls) using the measurement of interest (such as their expression levels for the gene being tested). Then we walk through the ranked list of samples, incrementing the enrichment score every time a case is encountered and decreasing it for every control. The increments and decrements are weighted by the absolute values of the Z-scores, therefore giving more weight to aberrations of larger scale. We are interested in the maximum cumulative enrichment score in this iterative process (for more details see Materials and Methods).

Additionally, we observed that under the null hypothesis with cases and controls uniformly ordered, the enrichment scores have a higher variance later down the walk (It can reach higher values by chance, given more steps). This can introduce a positional bias and decrease the power of the test. We show that this variance can be analytically computed without approximations. Consequently, we can correct for the positional bias by adding a standardization step for the enrichment scores at every position. The maximum standardized enrichment score is taken and its significance is assessed with permutations. Figure 1, shows an example of a standardized enrichment score computed using CRBN gene expression levels on Alzheimer disease data (Section 2.3.1).

**Figure 1.**
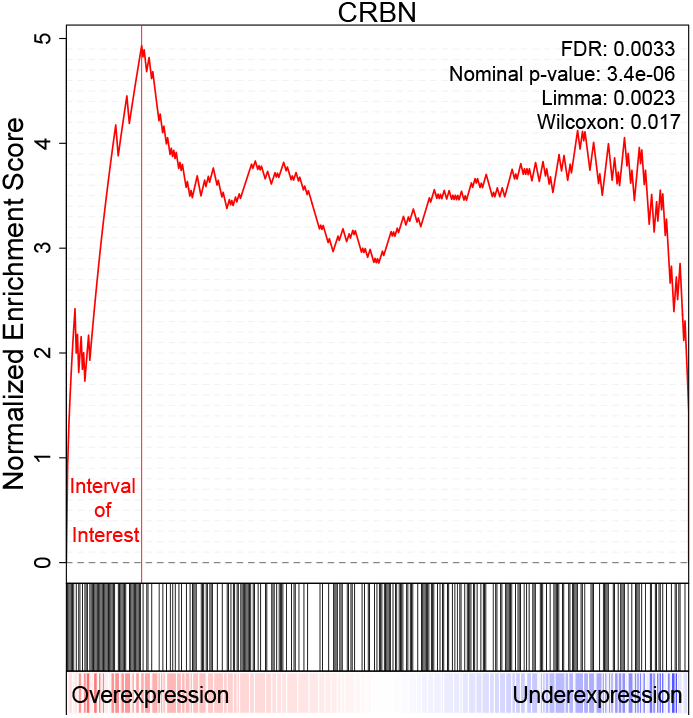
Example of a computed enrichment score. All individuals were ranked by decreasing expression of the *CRBN* gene. The enrichment score goes up whenever we encounter a case and goes down whenever we encounter a control. The maximum standardized score reached is 4.93 and corresponds to an uncorrected p-value of 3.4E – 06 and FDR of 0.003 for our test. There are 52 cases (19% of all cases) and 9 controls (4% of all controls) among the individuals to the left of the maximum. This was taken from the Alzheimer data in Section 2.3.1 and it corresponds to the *CRBN* expression distribution shown in Figure 7. Note that Limma and Wilcoxon do not detect this gene as significant when simultaneously testing all 25000 genes (Uncorrected p-values are respectively 0.002 and 0.01)

Note that our test is looking for an enrichment of aberrations in a proportion of cases as compared to controls. It is therefore not symmetric in terms of case/control labels. Different genes will be detected as associated with the considered condition if cases and controls are switched. This is not the problem in a setting where the focus is on identifying patterns associated with the cases. In a general two-group comparison we can run our test in both directions.

### 2.2 Simulations

For our initial set of simulations, we start from Gaussian simulated variables and then create an aberration enrichment pattern. We generate simulations by varying the following parameters:

- *n*: Sample size. Number of cases. We simulate *n* cases and *n* controls.
- *r:* Proportion of the cases with an aberration in the considered gene.
- *m,s*: Mean and the standard deviation of the initial simulated Gaussian variable.
- *d*: Multiplier controlling the magnitude of the aberration. The proportion *r* of affected individuals will have their average expression shifted *d* × *s* away from the rest of the cases and controls.

Simulations provide a controlled setting to assess the validity and power of our test in comparison to the widely used parametric and non-parametric approaches such as t-test, Wilcoxon, Levene and Kolmogorov-Smirnov test. The pattern of aberration enrichment simulated here could still be captured by traditional methods testing for a shift in mean or variance between cases and controls. We first simulate a variable with an aberration enrichment pattern of association by sampling the cases and controls from a Gaussian and then perturbing a proportion *r* of the cases. The perturbation is a shift by *d* times the standard deviation in one selected direction (an increase or a decrease). Figure 2 is an example of what the simulated data looks like given different parameter choices.

**Figure 2.**
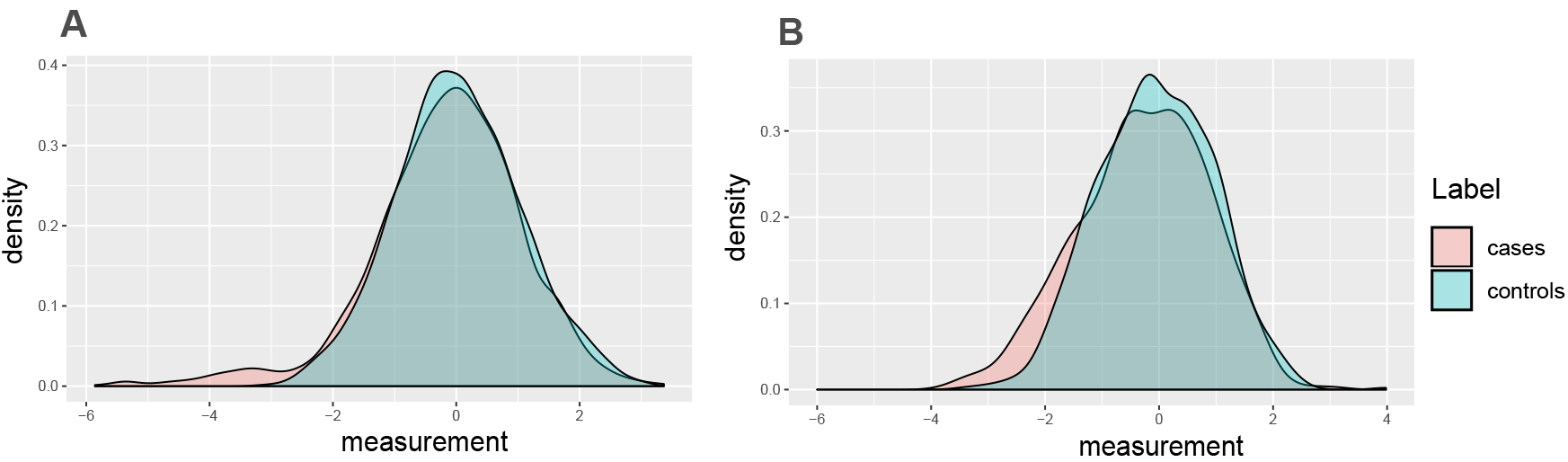
Examples of considered scenarios for aberration enrichment pattern. **A** *r* = 0.05, *d* =3, *n* = 500 **B** *r* = 0.15, *d* = 1.5, *n* = 500

We then tested the ability to detect these introduced aberration enrichment signals for our and the other parametric and non-parametric tests. Varying the sample sizes and the simulations parameters, we assess how often each method is able to detect the association given a nominal p-value threshold of 0.05 or a more stringent Bonferonni threshold of 2.10^−6^ (typically used in gene expression analyses to correct for multiple hypothesis testing). In these simulations we are generating one variable at a time and are assessing the power to detect that true association. Type 1 error is discussed in the next Section in a full realistic simulation setting with large numbers of associated and non-associated variables and again later on real data (For example in Figure 7-D). For every set of simulation parameters we repeat the experiment 200 times. The signal is considered detected if the p-value is less than the chosen threshold in more than half of the repeats.

#### 2.2.1 Power comparison

Figure 3 shows that variation in parameter space leads to three distinct outcomes.

**Figure 3.**
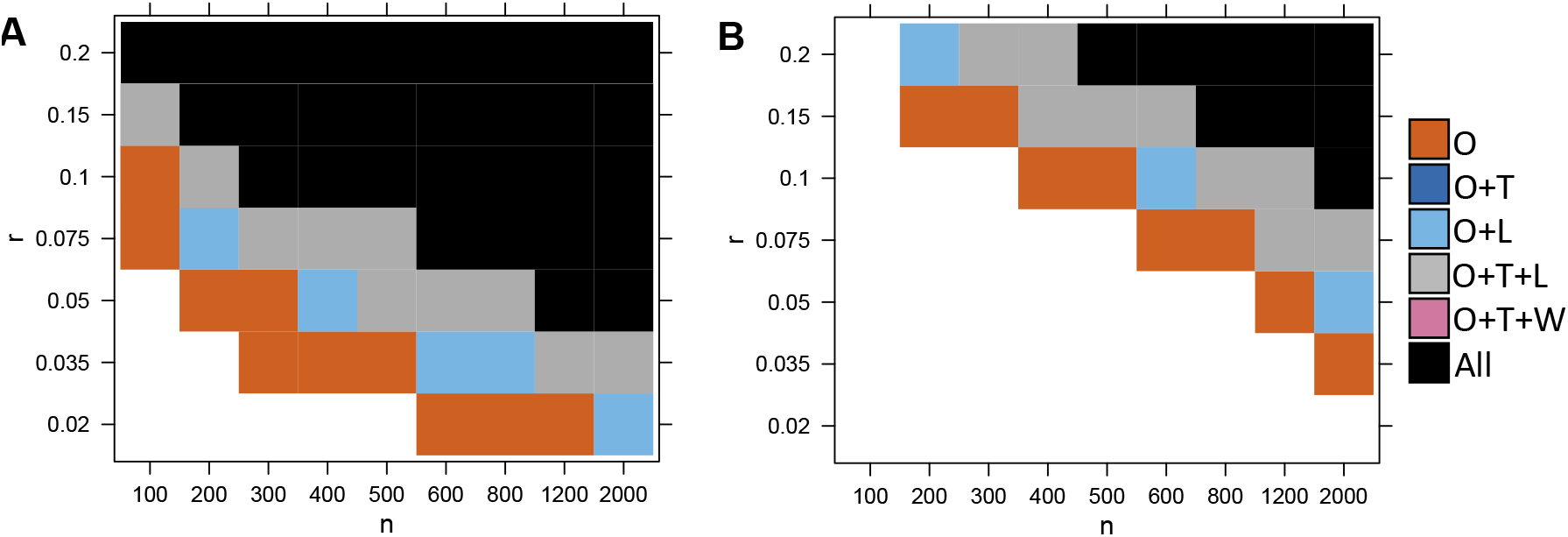
Ability of the different tests to detect the association, depending on simulations parameters *n* (sample size) and *r* (proportion of affected cases).In **A** a nominal p-value threshold of 0.05 is used. In **B**, a lower p-value threshold of 2.10^−6^ was used to mimic a realistic data analysis scenario where correction for multiple tests is required. We compare our test (O), the Levene test (L), t-test (T) and Wilcoxon (W). A method is able to detect the signal if the p-value is lower than the threshold in the the majority of 200 reruns. Here we show the results for *d* = 3. White is for the set of experiments where no method detected the signal, vermillion(red-orange) is when only our test detected the signal, light blue is when our test and the Levene test both detected it, gray is when our method, the t-test and the Levene test detected it and black is when all considered methods can detect the signal (including the Wilcoxon test).

- When either the proportion *r* or the sample size *n* is too small, no test can detect the association. The number of cases with an observable aberration is too low to generate enough statistical power (white region).
- When the proportion *r* and the sample size *n* are large, the number of cases with aberrations is high enough to create a significance shift in the mean (or variance) of the distribution of the cases compared to the distribution of the controls. In this case, most methods are able to detect the differential expression (black/ gray/ blue).
- Between these two regions, there is a domain where only our test is powerful enough to detect the association due to aberration enrichment (vermillion-red region).

Overall, we found that our test performs best, followed by the Levene test and the t-test. Wilcoxon had a lower power than the t-test and finally the Kolmogorov-Smirnov test was the least powerful (Kolmogorov-Smirnov test results not shown here for clarity purposes. See Supplementary Table 5). The Levene test performed slightly better on average than the t-test in this context (with *d* = 3). Figure 4 further shows there is a large difference in power between our test and the other tests in terms of magnitude of p-values, which was often several orders of magnitude lower for our test. This is especially true for the lower values of *r* (the signal is present in a smaller proportion of patients). For example, when *r* ≤ 0.05 the p-values returned by our test are often more than 4 order of magnitude (10000 times) smaller than those returned by any other methods.

**Figure 4.**
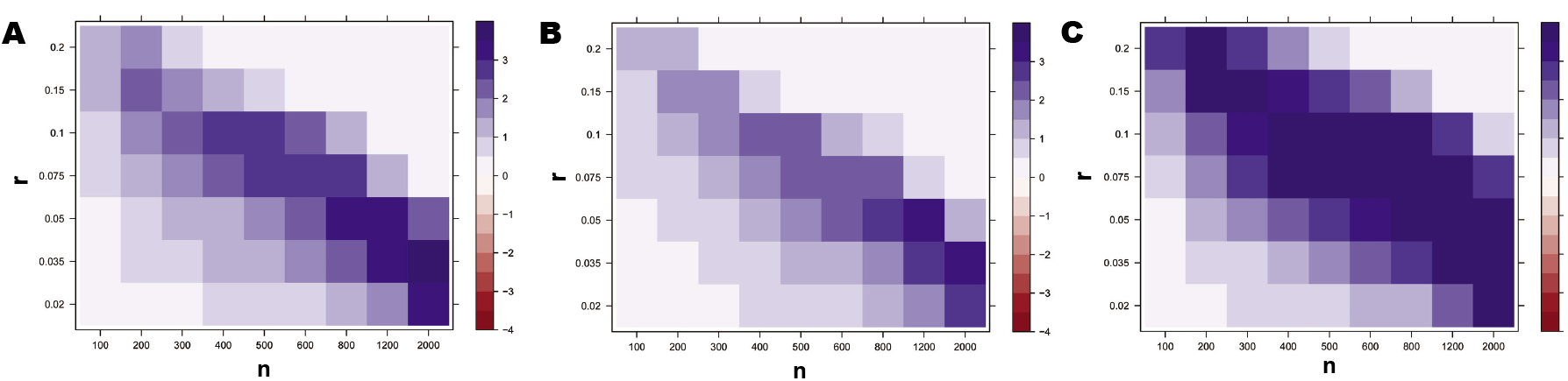
Comparison of the p-value magnitude between our test and **A t-test test, B Levene test, C Wilcoxon test**, depending on the simulation parameters *n* (sample size) and *r* (proportion of affected cases). Here we show the results for *d* = 3. The colors indicate the difference in log_10_ between the p-values returned by both method. For example 2 indicates that our test’s p-value is two orders of magnitude smaller (×10^−2^) than that of the Levine test. We capped the maximal difference at 4 for visual clarity. We ran 10^8^ permutations to compute the p-values for our test, therefore we set the minimal p-value to 10^−7^ for all methods in order to avoid artifacts of p-values estimation accuracy.

We also explored different scenarios by varying the other simulation parameters. We found that changing the mean *m* or variance *s* of the Gaussian had no effect on performance. In contrast, changing the value of the *d* parameter (perturbation magnitude multiplier) had a clear effect on performance in Figure 5. Higher values of *d* meant more cases became clear outliers for the expression of the considered gene, which makes the aberration enrichment signal easier to detect for our test. Higher *d* also means a higher effect on the overall mean/variance of the cases, therefore the power increases for all methods. Inversely, lower values of *d* negatively affect the performance of all methods. The ordering of the methods is overall maintained across the values of *d* with our test having the best performance in all scenarios followed by Levene/T-test and then Wilcoxon. However, we observe that the lower values of *d* are more severely affecting the Levene test compared to other methods. For *d* ≤ 2, the t-test start outperforming the Levene test (dark blue instead of light blue). This is expected because higher magnitude of perturbations have a larger effect on the variance.

**Figure 5.**
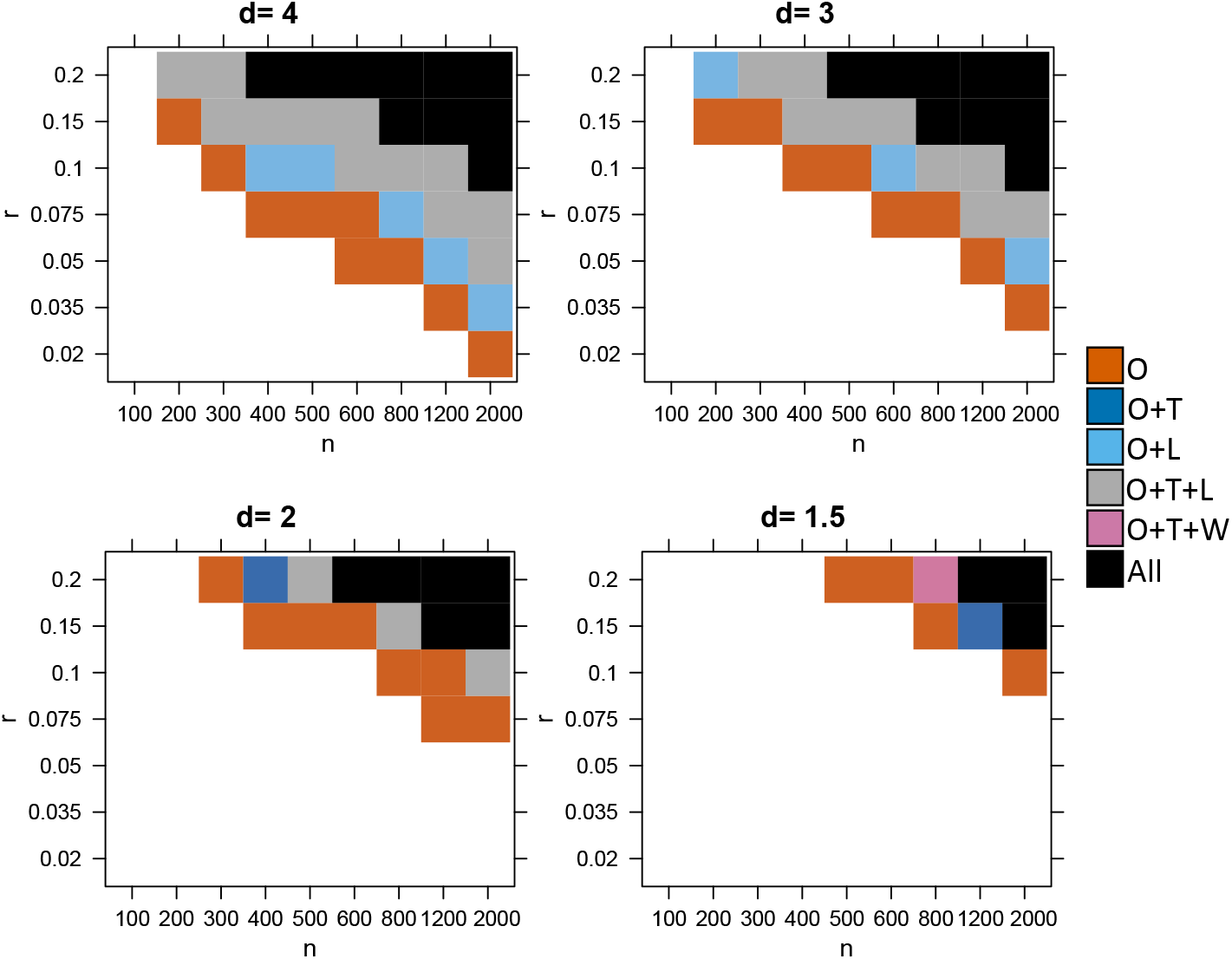
Ability of different tests to detect the simulated association, depending on simulations parameters *n* (sample size), *r* (proportion of affected cases) and *d*. p-value threshold of 2.10^−6^. A method is able to detect the signal if the p-value is lower than the threshold in the the majority of 200 reruns. We compare our test (O), the Levene test (L), t-test (T) and Wilcoxon (W).

Next, we studied the limitations of our test by increasing the values of *r*. For *r* = 1, there is no heterogeneity, i.e. all cases are affected the same way. While lower values of *r* correspond to the aberration enrichment (or heterogeneous response) scenario targeted by our test. Figure 6 shows that our test continues to be more powerful up to *r* ≤ 0.5 with a large difference in p-value magnitude for *r* ≤ 0.3. For larger values of *r* ≥ 0.7, there is no longer an advantage over using a t-test. We observed that the Levene test performance drops dramatically for the higher values of *r* and can no longer find the signal that is detectable by all other methods (pink region). For *r* = 1, we are essentially simulating cases that are mean shifted from the controls with no effect on the variance.

**Figure 6.**
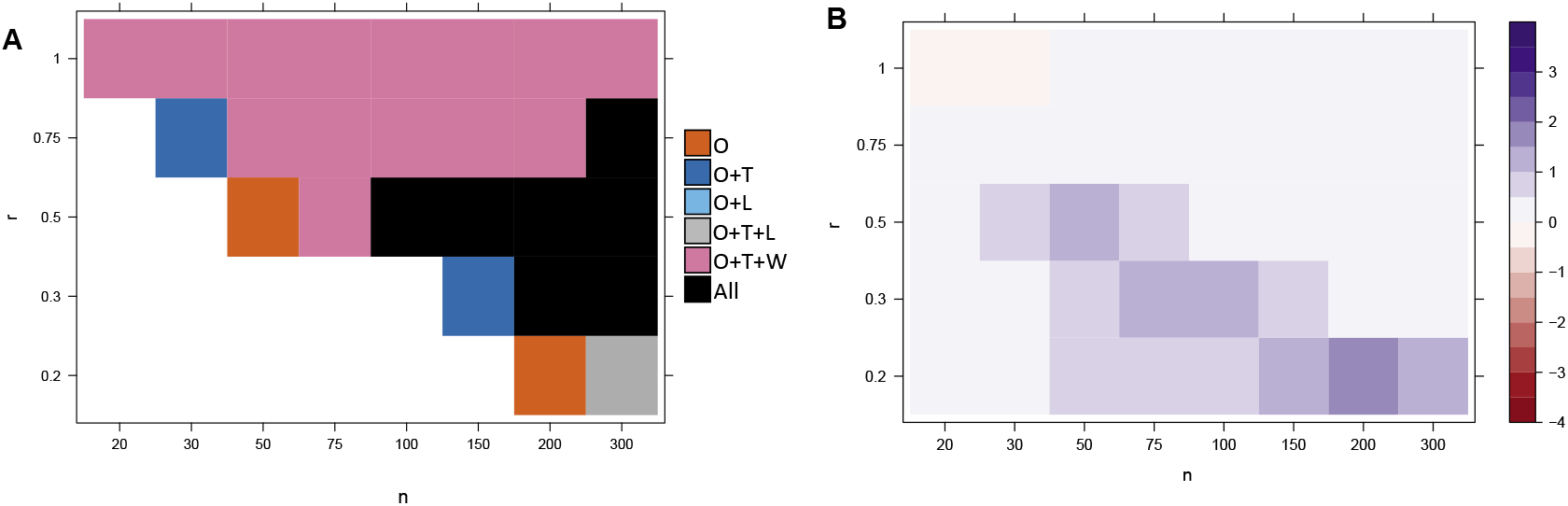
Results under less heterogeneity, i.e. higher values of *r* **A** Ability of the different tests to detect the association for higher values of *r*. A method is able to detect the signal if the p-value is lower than the threshold in the the majority of 200 reruns. Here we show the results for *d* = 3. We compare our test (O), the Levene test (L), t-test (T) and Wilcoxon (W). **B** Comparison of the p-value magnitude between our test and the best out of t-test, Levene and Wilcoxon test. depending on simulations parameters *n* (sample size) and *r* (proportion of affected cases). Here we show the results for *d* = 3. The colors indicate the difference in log10 between the p-values returned by both method. For example 2 indicates that our test’s p-value is two orders of magnitude (100 times) smaller than that of the Levine test. We capped the maximal difference at 4 for visual clarity.

Other scenarios: very low values of *r* (Supplementary Figure 1 and lower values of *d* ≤ 1 (Supplementary Figures 2 and 3), can be found in the supplementary materials. Overall, we conclude that our test can be a powerful alternative to currently used methods for scenarios with *d* ≥ 0.7 and *r* ≤ 0.5. Large gain in statistical power are obtained especially in settings with *d* ≥ 1.5 and *r* ≤ 0.3, meaning that less than 30% of the case group are different from the controls. We recommend using our test when heterogeneity is suspected.

#### 2.2.2 Simulating genome wide expression data

In the previous section, we showed that our test was more powerful than a t-test or a Levene test to detect the aberration enrichment pattern under the simplistic assumption of perturbed Gaussian when simulating a single variable at a time. In the differential expression literature, there are approaches that work on genome wide expression datasets (instead of a single gene) and that are more powerful than a t-test [25] for finding differentially expressed genes. Here we generated simulations of genome wide gene expression datasets based on real microarray expression data. The goal is to assess the performance of our test (power and type-1-error control) in comparison to other methods in the realistic setting typically considered when analyzing differential expression. By perturbing only a subset of all genes, we can also assess whether the different methods are well calibrated by analyzing the rate of false positives.

This setting also allows us to compare to the widely used method Limma [52] that is applicable to genome wide expression datasets. Limma uses empirical Bayes to borrows information across genes in order to empower the detection of differential expression, especially for lower sample sizes. Its efficiency has been proven in methods reviews publications where it always showed better or on par power and false positive control compared to all state-of-the-art methods, for both microarray [25] [44] and RNA-Seq experiments [61] [11].

Finally, we also compared our test to a joint test of scale and location by combining both Limma and Levene tests with the Fisher’s method as proposed in [60].

We use the 238 healthy controls from GSE63063 [62] from the Gene Expression Omnibus (GEO). We created our dataset by sampling gene expression from the real gene expression data and adding Gaussian noise *s* for a simulated *n* cases and *n* controls. We randomly selected a set of *g* genes as disease-associated amongst the 25549 genes in the data. The controls were left untouched but a proportion *r* of the cases were perturbed in every disease-associated gene, similarly to what we did in Section 2.2.1 (the expression values for the considered gene and the selected cases, were shifted in one direction by a factor of *d* times the standard deviation of the gene). We fixed the simulation parameters to *r* = 0.1, *d* = 3, *s* = 0.01, *g* =10 and varied the sample size n. We ran our method and the other tests (Limma, Levene, Combined test) on the full expression data for every choice of sample size and assessed the Type 1 error and the power of each method. More specifically, we measured the false positive rate using an FDR threshold of 0.1 and the true positive rate (proportion of the true genes that were detected).

Table 1 shows a difference in power similar to what we observed in the previous section, where we compared our test with the t-test and Levene test. In fact, applying the t-test to the full expression data results in p-values very similar to the ones generated by Limma. It seems that borrowing information across genes might not be helping Limma to noticeably improve performance over the t-test, for the sample sizes considered *n* ≥ 150 and in the setting of an aberration enrichment pattern. Additionally, we verified that using a logistic regression or ANOVA results in p-values that are equivalent to the t-test p-values in this setting (See Supplementary Table 5). We also verify that our test is more adapted to the detection of aberration enrichment in whole genome gene expression than Limma, t-test or the Levene test. We confirm our previous conclusion that when the signal is detectable, there is a considerable difference in power between our test and existing differential expression methods. This performance gap becomes wider for smaller values of *r* (proportion of affected cases) as shown in Supplementary Figure 4 where we repeated the same experiment with *r* = 0.05. This is expected given that there will be less of an impact on the mean or variance of the cases’ distribution when fewer individuals show aberrant levels for the gene of interest, therefore giving a larger advantage to our test since it does not rely on detectable broad differences between all cases and controls.

**Table 1.** Comparison of the false positive rate and the power of our test, Levene test, Limma and the combined test of scale and location. The average performance over 1000 simulations is shown here.

| n | 150 | 300 | 400 | 600 |
| --- | --- | --- | --- | --- |
| Limma/t-test FDR | 0.09 | 0.09 | 0.101 | 0.092 |
| Levene test FDR | 0.07 | 0.098 | 0.101 | 0.088 |
| Our test FDR | 0.1 | 0.087 | 0.084 | 0.082 |
| Limma/t-test power | 0.003 | 0.048 | 0.159 | 0.568 |
| Levene test power | 0.002 | 0.068 | 0.251 | 0.726 |
| Our test power | **0.006** | **0.658** | **0.908** | **0.992** |
| Combined test FDR | 0.394 | 0.317 | 0.301 | 0.303 |
| Combined test power | 0.034 | 0.552 | 0.853 | 0.992 |

Combining Limma and the Levene test in one joint scale-location test gives higher power than either test used separately as shown in Table 1. However, the joint test is not well calibrated in this setting as is illustrated by the high false positive rates limiting its applicability in a real data setting. Moreover, our test is still more powerful than the combined test, especially for lower values of r, as shown in Supplementary Figure 4. The only examples where our test was not significantly more powerful than the combined test in this setting with *r* = 0.1 is when the sample sizes were too low to detect the signal (*n* = 150) and when the sample sizes were too high (*n* = 600) and the associations were easier to detect. In comparison to all other methods such as the t-test, Limma, Levene, Wilcoxon, and the Kolmogorov-Smirnov test (See Supplementary Table 5), our test was significantly more powerful in all scenarios: p-values ≤ 2 × 10^−16^ in any comparison when *n* ≥ 300; and p-values ≤ 3 × 10^−3^ in any comparison when *n* = 150).

We conclude that our test is indeed well calibrated and that it is significantly more powerful than current tests to detect true associations when the signal takes the form of an aberration enrichment rather than a global shift in mean.

### 2.3 Results on real data across diseases

We downloaded, preprocessed and analysed case-control gene expression datasets from Gene Expression Omnibus (GEO). The sample sizes for each dataset are summarized in Supplementary table 1. The preprocessing involved removing 30 hidden (latent) factors with PEER [63] as was done in GTEX study on rare expression aberrations [33]. Since the simulation showed that Limma was as good or better than t-test, we ran Limma and our statistical test for each of our real data studies and analyzed the genes detected by each method. Our results on Wilcoxon test can be found in the Supplementary. As we can see in Table 2, there was a number of differentially expressed genes that were detected by both methods. In this analysis, we focus on the novel genes that were only found by our test. Additionally, whenever an association is found by our test, we can identify which subgroup of individuals and which interval of aberrant expression contributed to the test statistic and compute an estimated value of *r* (proportion of cases affected) for that gene (See Methods Section 3).

**Table 2.**
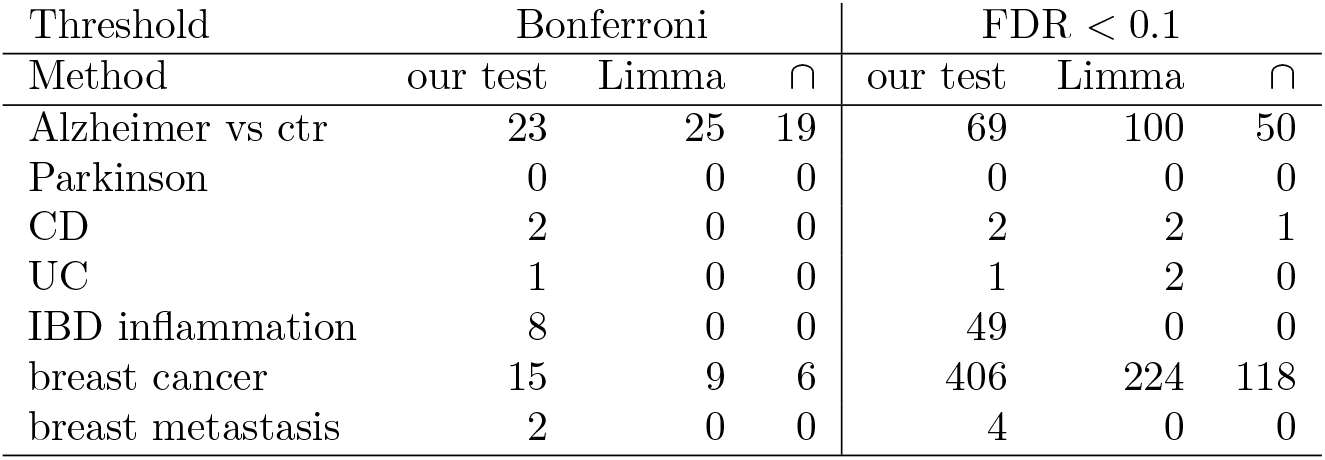
Real data findings: Number of genes detected

#### 2.3.1 Alzheimer and Parkinson disease

The GEO dataset GSE63063 [62] contains gene expression of 284 Alzheimer disease patients (AD), 189 mild cognitive impairment patients (MCI) and 238 healthy matched controls measured in blood. We ran Limma and our test to find genes that are differentially expressed between Alzheimer patients and healthy individuals. In this analysis we want to find novel genes that would not be picked up by Limma or genes that are much more significant by our test. Those genes would be over the diagonal in Figure 7-A. We observe that genes such as *UQCRH, ATP6V1D, CRBN, POMP, EIF3E* fit this criterion with the first two below Bonferroni significance threshold and the last three with FDR< 0.01. *UQCRH* is part of the KEGG pathway for Alzheimer disease, listed in both the organism specific and the conserved biosystems. *CRBN* or Cereblon (*FDR* = 0.003) is known to play a role in memory and learning and it has been previously associated with mental retardation [72]. It is also used in ubiquitination/proteasomal degradation of Tau [58] which could be relevant for Alzheimer disease [23]. *MYL6* only reaches significance by using our test (Bonferroni-corrected) while being under the significance threshold using Limma.

**Figure 7.**
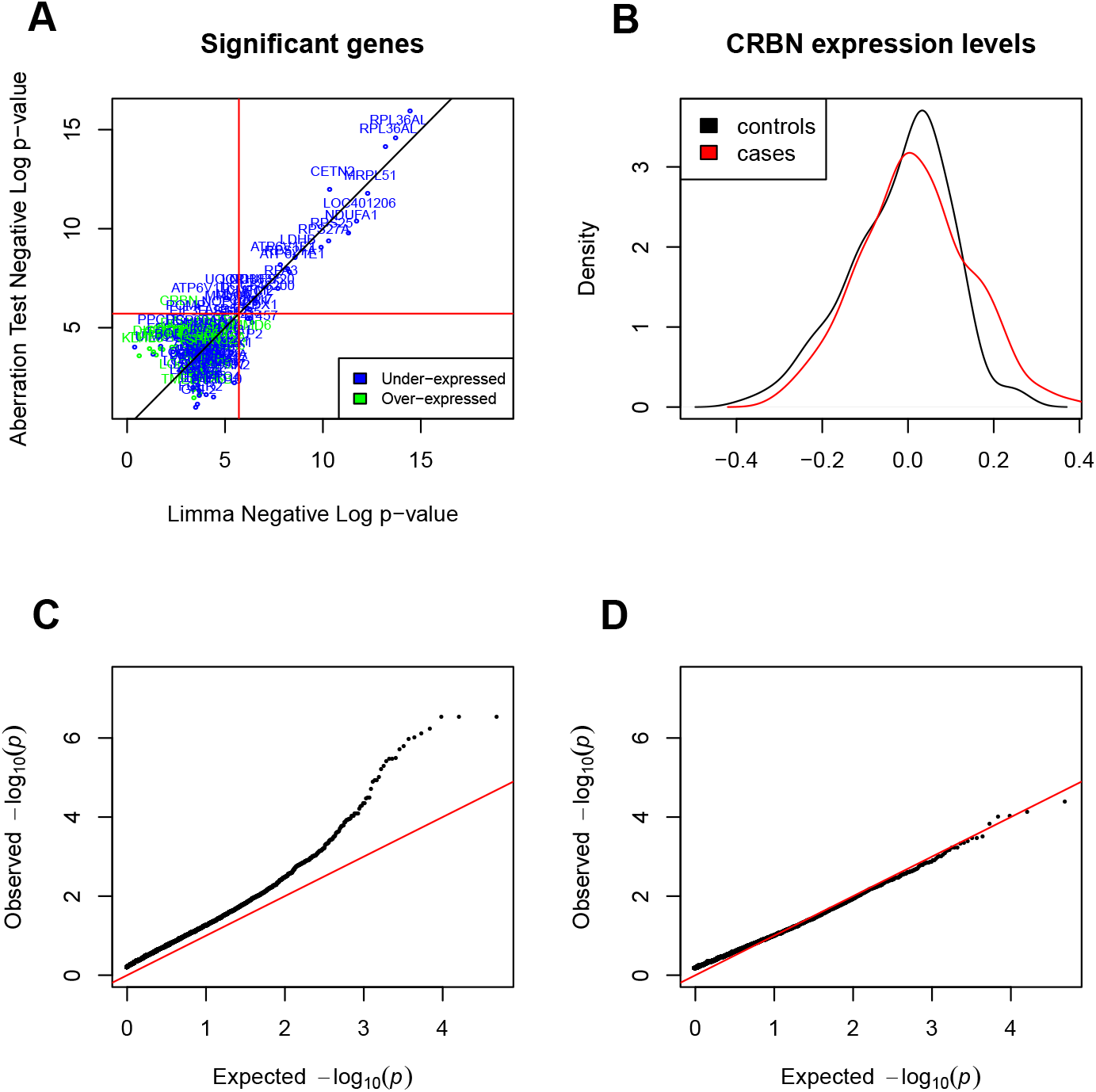
Results on Alzheimer’s disease. **A** Genes differentially expressed or aberration enriched in Alzheimer versus healthy controls, discovered using our test versus Limma. The genes that have *FDR* ≤ 0.1 for either method are plotted. Red lines correspond to the Bonferroni significance threshold. **B** Distribution of expression levels in cases and controls for the gene *CRBN* discovered by our test. **C** QQ plot of the p-values returned by our test on the Alzheimer data. **D** QQ plot of the p-values returned by our test after randomly permuting the samples.

The top 10 aberration enrichment genes (which exhibit a highly heterogeneous signal corresponding to an estimated *r* < 0.3), with *FDR* < 0.1 by our test, are: *CRBN* (ILMN1668582), *PPCDC, FBP1, DDX17, SYT13, GPER, DISC1, LRP3, TLR2, DNAJA1*. Even though the expression levels were measured in blood, many of the genes found are known for their functions in the brain. Furthermore, some of those genes have been shown to be involved in Alzheimer disease in the literature. For example, *DNAJA1* mediates Tau clearance [1] and *TLR2* is a major receptor for Alzheimer’s A*β* with a proven role in activating neuroinflammation [35]. These novel associations would not be detectable by other methods, showing the importance of going beyond differential expression and looking for heterogeneous effects and the aberration enrichment pattern.

The majority of associations uncovered in Figure 7-A are in the form of an under-expression of the considered gene in Alzheimer patients. Exceptions to this are *CRBN, DDX17, GPER, DISCI, LRP3* and *TLR2*.

On Figure 7-B, we plotted the distribution of *CRBN* expression levels in cases and controls to better show the nature of the association, where a subgroup of cases have aberrant overexpression of the gene. Figure 7-C shows that there could be a large number of genes exhibiting some aberration enrichment signal in association with Alzheimer. Comparing the QQ plot in Figure 7-C to the one in Figure 7-D where we permuted the labels confirms that the associations discovered are not spurious hits due to a badly calibrated statistical test but signals that are truly associated with the case-control labels.

We performed a similar analysis of Parkinson disease (IPD) and found no associated genes using any of the considered methods. The dataset (GSE99039 [56]) consisted of whole blood gene expression data for 205 IPD cases and 233 controls.

#### 2.3.2 Inflammatory Bowel Disease

The inflammatory bowel disease data (GSE73094 [47]), contains the gene expression of 712 preselected genes, including 440 genes in IBD GWAS risk loci and 15 housekeeping genes, in 608 samples from Crohn’s disease (CD) patients, 331 from Ulcerative Colitis (UC) patients and 50 samples from non-IBD individuals. The samples were taken from the colon and terminal ileum. Overall 374 of the samples were taken during inflammation and 609 taken from non-inflamed tissues (6 samples with missing inflammation status were removed).

We first looked for genes associated with CD versus UC and vice-versa by considering the not inflamed samples which were more numerous than the inflamed ones. This resulted in 181 UC and 314 CD non-inflamed samples. After preprocessing (See Methods Section), we looked for genes associated with CD and genes associated with UC by comparing each group to the other.

Only one gene was significantly associated with UC. *C11orf9* was detected by our test. The gene is also called *MYRF* and was previously mentioned in the IBD literature as part of a co-expression cluster of upregulated genes [78] and the nearby SNP rs4246215 was previously associated with IBD in GWAS [26,39].

Two genes were found to be significant for CD: *BTNL2* and *IRF4*. Both were detected as significant only with our test. *IRFR* was discovered by Limma too with *FDR* = 0.09 (*FDR* = 6*E*–4 by our test). *BNTL2* was not a broad effect (*r* = 0.15) and therefore it was not detected by other methods.

We also looked for genes associated to inflammation status across all conditions by taking all samples from the original data (374 inflamed and 609 non inflamed) and correcting for disease type as a confounder.

In figure 8, 8 genes were found to be significantly associated with inflammation. *PLF4, ITLN1, IL24, S26A3, PIGR, RNT2, SL9A4* and *FAM55D*. All of them were only detected with our test.

**Figure 8.**
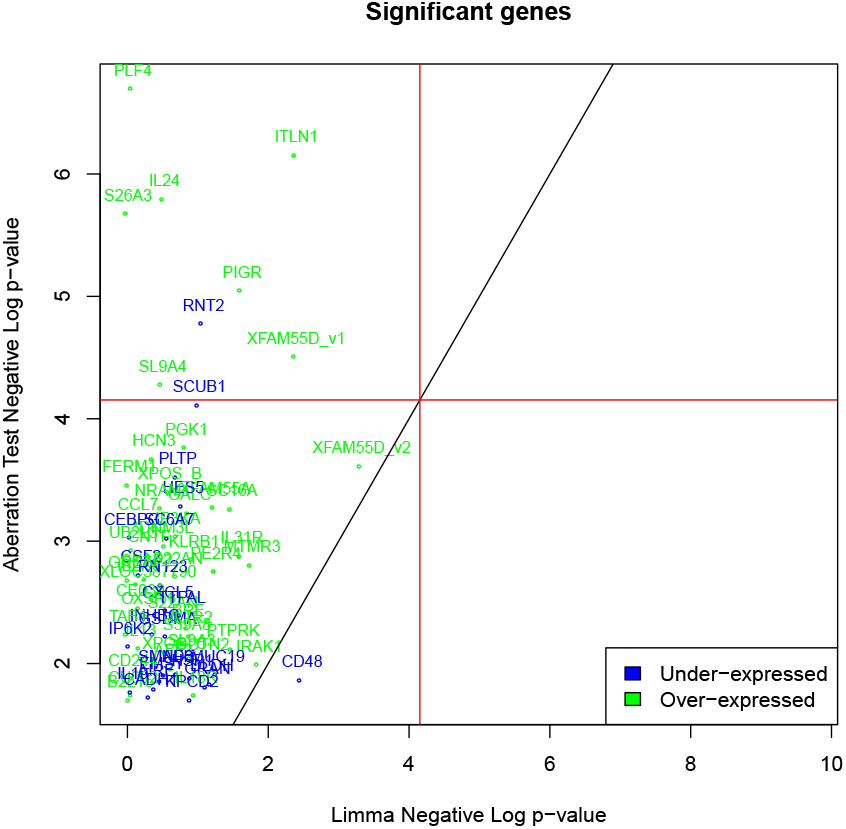
Gene expression sssociations in IBD, inflamed versus non-inflamed samples, discovered using our test versus Limma. The genes that have FDR> 0.2 for either method are plotted. Red lines correspond to the Bonferroni significance threshold.

#### 2.3.3 Breast Cancer

The heritable breast cancer data (GSE47862 [50]) contains gene expression in peripheral blood in 158 women with heritable breast cancer and 163 healthy controls. The top 3 associated genes by our test were Entrez-id 100129342, *PIK3C2B* and *CR2*. We identified a large number of associated genes with heterogeneous effects (estimated *r* < 0.3). The top 10 genes of our gene list are: *PSIP1, SLCO2B1, TLX3, CDKAL1, MCMDC2, GPAA1, B4GALT1, FUT4, PIGR, CDCA7* (*FDR* < 0.025). Several of these genes have substantial evidence in the literature of their involvement in breast cancer. For example, it is known that silencing *CDCA7* in triple negative breast cancers reduced tumorigenicity and invasion in [74], while the forced expression of *GPAA1* in [71] was shown to increase them. PSIP1 has also been shown to directly promote tumorigenicity in breast cancer [59]. The expression of *SLCO2B1* was shown to be significantly correlated with histological grade in ER+ breast cancer [38]. *TLX3* is also known as T Cell Leukemia Homeobox 3 and is a transcription factor oncogene. rs9368197 in an intron of *CDKAL1* is associated with breast cancer risk [8]. Knock-down of the *B4GALT1* gene and the inhibition of its function has been shown to inhibit the estrogen-induced proliferation of breast cancer cell lines [9]. *FUT4* has been proposed as an effective diagnosis biomarker of breast cancer [73]. *PIGR* is known to be upregulated in breast cancer, and other cancers [69].

Given the large number of associations we run a gene set enrichment analysis on Reactome [17]. The top module was TP53 regulation of metabolic genes. It was not significant after correcting for multiple hypothesis.

If we consider only the very heterogeneous effects (estimated *r* < 0.1), we find the following associations with FDR < 0.1: *CMKLR1, CASC5*(*KNL1*), *SUSD1, RASSF4, AOC’4P, ADHFE1, FAM71C, RRN3P3. CASC5*(*KNL1*) is known as Cancer Susceptibility Candidate Gene 5 Protein. *RASSF4* is a member of the *RASSF* family of tumor suppressors. *AOC4P* is a lncRNA involved in Hepatocellular Carcinoma and Colorectal Cancer [22] and *ADHFE1* is a breast cancer oncogene [41].

The breast cancer metastasis data (GSE48091 [37]), measures the gene expression in primary breast cancer tissue in 166 cases where metastasis happened and 340 cases without metastasis. Only 2 genes reached the Bonferroni significance threshold for association with metastasis status: *TAOK1* and *BC04 2012*, and 2 more had low FDR: *RALB* and *CA428624* and they were only significant with our test (*FDR* = 0.02). Both *TAOK1* and *RALB* were found to be underexpressed. *TAOK1* had a relatively low p-value by Limma (E-04) but the 3 other associations are specific to our test and correspond to low values of *r*. *TAOK1* was previously listed as a metastasis-associated genes in basal-like breast tumors [18] (through expression). *RALB*, also known as RAS Like Proto-Oncogene B, which is known for its role in invasion and metastasis across cancers and specifically for breast cancer [76].

For both breast cancer datasets, we found interesting heterogeneous associations that would not be detected by tests looking for the broad differences between all cases and controls, illustrating the value in looking for aberration enrichment.

### 2.4 Results across other types of -omics data

We ran our test on publicly available miRNA expression and DNA methylation datasets that we downloaded from GEO. Given the larger size of these datasets (thousands of samples in miRNA datasets and more than 450k features for methylation data), we used *k* = 100 for the number of hidden confounders removed (using the PEER correction). Table 3 summarizes our findings with *k* = 100. The results with *k* = 30 are presented in the Supplementary Table 3.

**Table 3.** Real data findings: Number of genes detected

| Threshold | Bonferroni |  |  | FDR < 0.1 |  |  |
| --- | --- | --- | --- | --- | --- | --- |
| Method | our test | Limma | ∩ | our test | Limma | ∩ |
| RA methylation | 119 | 2 | 1 | 506 | 3 | 3 |
| Schizo. cortex | 22 | 0 | 0 | 139 | 0 | 0 |
| Schizo. blood | 75 | 0 | 0 | 530 | 0 | 0 |
| breast miRNA | 427 | 0 | 0 | 564 | 0 | 0 |
| ovarian miRNA | 364 | 5 | 4 | 849 | 23 | 23 |

Our test returned a larger number of associations in these datasets, most of which were not discovered by Limma (See Table 3). In fact, using Limma or a t-test resulted in few to no associations, especially in methylation data. We did observe bimodality or mutimodality in the methylation levels of many sites and that can partially explain the poor performance of t-test and Limma. Nonparametric methods such as our test and Wilcoxon might be better suited for methylation data. Wilcoxon test returned a larger number of associations compared to Limma and t-test (See Supplementary Table 4). These associations were still fewer in numbers than our test, and partially overlapped with our associations that corresponded to broad effects (large values of *r*).

In our analysis, given the large number of associations by our test, we prioritize those that affect a small proportion of cases (lower estimated values of *r*) rather than broad signals, these were also not found by Wilcoxon/Limma/t-test. Additionally, the large number of associations allows us to perform gene set enrichment analysis for the associated probes that are annotated to genes. Note that some of these associated probes are co-located and annotated with the same gene name. However, most of the genes we discuss below only have one associated probe.

In each dataset, to verify that our test is still well calibrated we performed random label permutations and showed that we find no associations under the null.

#### 2.4.1 DNA methylation in Rheumatoid Arthritis

The Rheumatoid Arthritis methylation data (GSE42861 [36]) measures DNA methylation for 354 RA patients and 335 controls. A large number of sites was found to be associated by our test. The top associations (FDR< 10^−4^, *r* < 30%) involve multiple sites near HLA genes such as *HLA-DQA1, HLA-DPB2, HLA-DRB1, HLA-DMB*, along with other genes: *SLC43A2, MBD1/CXXC1, ALLC, LPP, ESYT2, NMB*. We also find other HLA genes such as *HLA-B, HLA-DRB5, HLA-DQB1, HLA-DRB6* that are associated with *FDR* < 0.01. *HLA-DRB1* is the strongest causal gene for RA [67]. In our data, 53 methylation sites were annotated to *HLA-DRBl*. Among these, 4 were found to be strongly associated by our test exclusively: cg04026937 (*FDR* = 1.3e – 03), cg06204447 (FDR = 8.1e – 05), cg18052547 (FDR = 1.4e – 02), cg23899527 (FDR = 2.2e – 05). One (cg00598125) was found to have some association by Wilcoxon (*FDR* = 0.06) and was subsignificant by our test (*FDR* = 0.19) and did not correspond to an aberration enrichment pattern of association (r = 0.6). We also looked at loci that are dysregulated in less than 10% of patients with FDR< 0.1. Among the 12 loci we find *CHI3L1* (hypomethylated in *r* = 9%) which is a rheumatoid arthritis autoantigen [10], SNW1 (hypomethylated in *r* = 7%) which is a nuclear factor kappa B (NF) regulatory gene involved in RA pathogenesis [53] and *CAV1* (hypermethylated in *r* = 9%) which is involved in NF-kappa-B activation in a T-cell receptor/CD3-dependent manner [45]. Among the genes mentioned above, *HLA-DQA1, HLA-DQB1, SLC%3A2, MBD1/CXXC1, NMB* and one site near each of *HLA-DRB1*, are associated by Wilcoxon with FDR< 0.1. These common associations always correspond to the higher values of *r* (*r* ∈ [0.26 – 0.3] when we selected only the hits with *r* ≤ 0.3). All other associations are uniquely found by our test and many of which correspond to lower values of *r* ≤ 0.2.

An enrichment analysis on Reactome [17] shows several immune system modules are enriched in the candidates returned by our test with *r* < 0.3 and FDR < 0.1. The module Class I MHC mediated antigen processing and presentation is strongly enriched (FDR < 8.10^−15^). The responsible genes are *CLCN7, TRIM41, ERICH1, HLA-B, TAP1, UBE2E2, UBAP2L, CBLB, TAPBP*. The corresponding submodules: Endosomal/Vacuolar pathway, ER-Phagosome pathway and Antigen Presentation: Folding, assembly and peptide loading of class I MHC, are particularly enriched (FDR < 8.10^−15^). The module Interferon Alpha/Beta signaling is also enriched because of genes *ZNF605, HLA-B, TAP1* (FDR < 8.10^−15^).

While other methods are unable to recover the previous modules, the associations with Wilcoxon agree with our test on other modules containing *HLA-DRB5, HLA-DQA1, HLA-DRB1, HLA-DQB1*: TCA signaling, PD-1 signaling, Interferon Gamma signaling, MHC class II antigen presentation (the latter also containing *HLA-DMB, CAPZB, TAP1, HLA-DOB*). This is not surprising since MHC class II antigen presentation is very well known to be involved in RA [65]. Finally, there are several modules related to NOTCH signaling which is also known to play a role in RA [46] (containing the following associated genes *NCOR2, HDAC4, HDAC2, SNW1, ERICH1, FBXW7, MIB2, PSEN2, RBPJ, NOTCH4*). Overall, our test detected several associations with disease-relevant genes and pathways, some of which were not detected using any other approach.

#### 2.4.2 DNA methylation in Schizophrenia

The schizophrenia methylation datasets (GSE74193 [24] and GSE80417 [21]) respectively describe the DNA methylation in dorsolateral prefrontal cortex and whole blood. After preprocessing (See Methods Section), the first dataset had 191 schizophrenia cases and 335 controls and the second dataset had 305 cases and 333 controls.

In the brain, the top loci with proportion of affected cases *r* < 30% are by the genes *HLA-DRB6, SOX2OT/SOX2*, intergenic region at loci cg23330385, *NAALADL2, LIN7A, SOAT1, HLA-DRB1, ALDH3B2, LOC81691, NMNAT2, CNRIP1, TTC23L* and *SLC16A12* (all associations are with FDR< 0.01).

Several of these genes have been implicated with schizophrenia in the literature. For example, *LIN7A* is at the overlap of several rare CNVs associated with schizophrenia in [12] and induced overexpression of *CNRIP1* is known to cause a schizophrenia like-phenotype in mice [48]. *NMNAT2* is important for the maintenance of neurons and is known to be neuroprotective in several models of neurological disorders [2], while *HLA-DRB1* is the most frequently reported genetic association to Schizophrenia [70]. Furthermore, looking at the sites with lower proportion of affected samples (*r* < 10%), we find the 6 associated sites with FDR< 0.1: *CYFIP1, ST6GALNAC1, ABCA8, CPSF6, C6orf25* and intergenic site cg25008182. The site near the *ST6GALNAC1* gene is hypomethylated in 7% of the cases, and is known to be associated (through hypomethylation) with schizophrenia and bipolar disorder in an identical twin methylation study who are discordant for these diseases [15]. *CYFIP1*, here hypermethylated in 8% of the cases, was previously associated with schizophrenia and Autism through CNVs and is known to regulate the balance between synaptic excitation and inhibition [14]. The *ABCA8* gene is important for lipid metabolism in oligodendrocytes, myelin formation and maintenance, and *ABCA13* from the same subfamily is associated to schizophrenia through GWAS [29]. The genes listed above are uniquely found by our test except for *NMNAT2* which was found with FDR= 0.08 by Wilcoxon and FDR= 0.002 by our test. This shows that looking for aberration enrichment in addition to traditional approaches can lead to novel associations that might improve our understanding of disease.

The Reactome gene set analysis found an overlapping group of gene sets previously found by Wilcoxon and our test of Rheumatoid Arthritis, such modules containing the genes *HLA-DRB5*, *HLA-DQA2*, *HLA-DRB1*: TCA signaling, PD-1 signaling, Interferon Gamma signaling, MHC class II antigen presentation (the latter also containing *RACGAP1* and *ITFG1*). In schizophrenia, these modules are only detected through our test and not through Wilcoxon. This result is not surprising and is consistent with the strong associations between the HLA locus and schizophrenia found in different studies [42]. We also report the following modules of unknown relevance to schizophrenia: Glucuronidation with FDR=8.81E – 04 (genes *UGT1A3* to *UGT1A10*) and Phase II - Conjugation of compounds with FDR=5E – 03 (*SLC35B3, GGT7, MGST3*, and *UGT1A3* to *UGT1A10*).

In blood, the results were less interesting with a very large number of associated sites in Table 3(Wilcoxon also found 61 and 242 sites by Bonferroni and FDR respectively) and less obvious associations with schizophrenia in previous literature among our immediate top genes. The top 10 associations with *r* < 0.1 that are close to genes, are near *AP2S1, MYH7, DSCR3, C14orf182, TMCO1, PRR25, LOC389333, SELS, XKR6, DGKZ*. All of these associations have FDR< 0.05. *DSCR3*: Down Syndrome Critical Region Gene 3 has previously been associated with neuroticism in a genome wide linkage study [4]. *C14orf182* has been associated with schizophrenia in a Whole Genome Sequencing study done in discordant twins [66]. *DGKZ* is located within a schizophrenia GWAS loci and is further known to be dysregulated in schizophrenia patients [3, 49].

Both in brain and in blood, our test is recovering novel associations with genes/loci potentially relevant to Schizophrenia which would not be picked by other methods because of the heterogeneous nature of these associations (aberration enrichment).

#### 2.4.3 miRNA in breast and ovarian cancer

The breast cancer miRNA data (GSE73002 [57]) describes the serum miRNA levels of 1280 breast cancer cases and 2686 non-cancer controls. The ovarian cancer miRNA data (GSE106817 [75]) describes the serum miRNA levels of 399 ovarian cancer cases and 3647 non-ovarian cancer controls (includes 859 samples from other cancers). After preprocessing (See Methods Section), we ran Limma and our test on both datasets. Overall 963 and 2565 miRNAs measurements were made in the breast cancer and ovarian cancer dataset respectively. Out of those measured miRNAs, a relatively large proportion was found to be associated to the cancer status according to our test as shown in Table 3. We attempted to use larger values for the number of PEER factors *k* but this did not substantially reduce the number of associated hits (Supplementary Table 2). For example, in the breast cancer dataset, our test uncovered 483 associated miRNAs for *k* = 30. Using *k* = 100 or *k* = 200 only reduced that number to 427 and 425 respectively. Similarly in the ovarian cancer dataset, 462 associations were detected by our test for *k* = 30 and that number reduced to 364 and 352 for *k* = 100 and *k* = 200 respectively. Using Limma or a t-test returned very few to no associations while Wilcoxon returned a smaller number of associations than our test.

To show that these associations are not artifacts from our test, we performed random permutations of the labels and found zero associations. Meaning that there does not seem to be an inflation for type 1 errors for our test.

One possible explanation of these results is that cancer generates a large number of effects that are not homogeneous across patients. This is a well known phenomenon [54]. Heterogeneous downstream effects of cancer might include events such as large copy number changes, structural variants, large effects on chromatin conformation and epigenetics. Any of these events can result in dysregulation of miRNAs and any single event could be happening in a smaller proportion of cancer cases. The heterogeneity of cancer presentations across patients could also lead to a heterogeneity of downstream effects that would be observed as a large number of associations by our test. This result is consistent with the large number of associations we also observed in gene expression data in the breast cancer dataset compared to non-cancerous diseases. In Section 2.3.3, we observed 453 genes with FDR< 0.1 and 1506 genes with FDR< 0.2 in association with breast cancer.

Under this assumption of numerous heterogeneous downstream effects, it is difficult to pinpoint miRNA dysregulations that would be drivers of cancer among a very large number of associations. This is particularly problematic when we have a high proportion of associated features among all features (> 20% of miRNAs have *FDR* < 0.1 in our data). This shows the limitations of directly applying our test to cancer, where there is an accumulation of heterogeneous passenger events.

However, we argue that our test can be used in this context, but not for the task of feature selection (identifying relevant cancer miRNAs). Instead, we use it for identifying features (in this case, miRNAs) that are helpful for classifying individuals into cases vs controls. The argument here is that even if (most of) the associations are just downstream heterogeneous effects, they can still be used as biomarkers of cancer.

For each dataset, we split the data into a discovery cohort and a held-out cohort (not to be used for feature selection or training). We run our test on the discovery data to uncover miRNAs with heterogeneous associations. Many of our associations are found with *r* < 30%, meaning the considered miRNA’s association is produced by only a proportion of individuals with extreme values (overexpressed or underexpressed). We define the intervals of expression that are responsible for the association (using the index at which the standardized enrichment score is maximal, See Figure 1 as an example), then we assign a value of zero to all other individuals that are not in the interval of interest. This manually introduced non-linearity helps the model focus on meaningful dysregulations rather than considering the full expression spectrum as a whole for each miRNA. We use a lasso-penalized logistic regression classifier (R package glmnet [20]).

We report our results in Table 4 where we used a combination of feature selection and a logistic regression classifier to differentiate cancer patients and healthy controls. We select either the top 300 features by Limma or the top 300 heterogeneous features (*r* < 0.3) by our test. We optionally transform the top features of our test by assigning a value of zero to all individuals outside of the interval of expression that drove the association (Top Hetero. transformed column). We also compare to using all features or only the features previously used in the literature for this classification problem (a set of 5 and 10 miRNAs respectively for the breast cancer data and the ovarian cancer data).

**Table 4.** Cancer-control classification performance on held-out data with Area under the Precision-Recall curve (AUPRC) after feature selection by different methods.

| Features | Top 300<br>Limma<br>features | Top 300<br>Heterogeneous<br>features | Top Het.<br>features<br>transformed | All<br>features | Literature<br>features |
| --- | --- | --- | --- | --- | --- |
| Ovarian Cancer | 0.695 | 0.743 | **0.948** | 0.696 | 0.503 |
| Breast Cancer | 0.530 | 0.537 | **0.965** | 0.632 | 0.541 |

Using the features selected by our test leads to a much better classification performance compared to when we use the top features returned by Limma or when we use all features in the classifier. Our approach reaches AUC and AUPR over 0.94 for both datasets which is a much better performance than other feature selection approaches such as using Limma. The transformation of keeping only the expression values within the aberrant interval defined by our test is very helpful. This is consistent with our previous observation in simulation experiments that a logistic regression is not good at handling/detecting features with heterogeneous effects 2.2.2. The non-linear data transformation based on our test results seems to address this limitation of logistic regression. In fact, using a non-linear classifier such as random forest (R package RandomForest [34]) leads to a very similar performance to using transformed features in the logistic regression case, but our approach is easier to interpret. The Random Forest AUPR is 0.92 and 0.98 on the held-out data respectively for the ovarian cancer and the breast cancer datasets (See Table 4). However, the random forest performance is unchanged (±0.01 AUPR) whether we use Limma’s top features or those of our test and whether we transform the features or not.

It is very important to note that the problem of classifying cancer cases from controls has already been solved with very high accuracy for the same miRNA datasets [57,75]. A full classification performance is achievable even with few features because of the very broad difference observed between data originating from cases and controls. For example, running Limma, Wilcoxon or our test on the non-preprocessed data results in almost all miRNAs being strongly differentially expressed. In this proof of concept, we used processed data where 100 hidden PEER factors were removed. Some of these factors correspond to broad signal of cancer that could easily separate cases and controls. In fact we verify that using 20 of those hidden factors as features, we can recover a perfect classification with logistic regression or random forest. By removing the 100 hidden factors from the data, we made the classification problem harder than the one previously solved on the original data. In this proof of principle, our goal is to prove that heterogeneous disease signals do exist and that they have predictive value beyond broad signal. Using our test to detect and process these heterogeneous signals, we showed that we can improve upon the performance of a linear classifier in an interpretable way.

## 3 Discussion

In this paper, we presented a statistical test for detecting a pattern of association different from an overall shift in mean or variance between cases and controls. We call this pattern “aberration enrichment” or association with “heterogeneous effects”. Our test works in a case/control setting with a continuous input variable (such as a gene’s expression) and scales to hundreds of thousands of variables.

Through the use of simulations, we showed that our test is more adapted at uncovering associations with heterogeneous effects compared to the widely used statistical methods. Our test is well calibrated and uses permutations to assess the significance of the results. The power of our test is inferior or on par to traditional approaches in the classical setting, i.e. for detecting broad signal with no heterogeneous effects, but becomes vastly superior when the signal of interest concerns a smaller proportion of the cases (*r* ≤ 30%).

By applying our test to complex diseases and several real gene expression datasets, we showcase its ability to detect novel disease-relevant genes that would not be detected by traditional differential expression methods. We further applied our test to other omics data types (miRNA and methylation) and reported novel associations.

Many of the genes found by our test do not exhibit a broad signal across the disease cases. This makes their association with the disease less likely to be a homogeneous downstream consequence of the disease itself. However, that does not imply these genes are causal for the disease. It is still possible that some confounding variables (such as the environment or a drug) is affecting a subset of the cases. It is also possible that the considered disease is heterogeneous enough to generate a multitude of heterogeneous downstream effects on the measurements that are unobserved in the controls. For example, cancer may generate heterogeneous downstream consequences such as large CNVs and chromosomal rearrangements which would appear to our test as consistent outliers enriched in cancer cases but not in controls. Our test cannot distinguish causal factors from heterogeneous downstream consequences. Similarly to the widely used differential expression approaches, our test can return a very large number of associations in some contexts, thus rendering a downstream search of causal elements very difficult.

In real data experiments, it is important to correct for known and hidden confounders in order to remove broad irrelevant signals and obtain a small set of associated genes. Here we used PEER [63] to correct for confounders. It is always possible that some complex/non-linear hidden confounders or other broad effects (such as cell-type proportion heterogeneity across patients) are not being fully removed by PEER. It is also possible that this procedure of removing hidden confounders might also be removing signals that are relevant to the causal mechanisms of the disease in our experiments (leading to false negative genes). Furthermore, the procedure of correcting for confounders generally works under the assumption that the confounders affect the mean, but some confounders could be affecting the variance of the measurement of interest [16]. If that is the case, it is possible to identify false associations driven by confounders that have different variances in cases and controls. Better upstream procedures for correction which consider the effect of confounders on variance will be beneficial for all the methods considered but especially so for methods that look for beyond the effect on the mean.

Currently, the need for permutations makes the method slower than the widely used statistical tests. Especially if we want to accurately measure very low p-values. More work needs to be done to better model our test statistic (the max over correlated standardized enrichment score variables) in order to obtain a closed form solution. Currently the null distribution over test statistics is not analytically computed and it does not clearly fit any known parametric distribution we tried. (The max over dependant standardized weighted hypergeometric variables is not easy to model. A polynomial approximation works to fit the tail but it is hard to justify so we did not rely on it).

The statistical test presented in this paper could be applied to other datasets and other fields beyond complex diseases and omics data. Wherever a 2-group test is used (such as Wilcoxon, t-test or the equivalent logistic regression), our test could be a complimentary analysis, especially where we might expect a non-homogeneous difference between the groups. For example, in randomized clinical trials, we often compare a continuous measure of response (a change from baseline in a measure of disease severity) between individuals who took the drug and individuals who took a placebo in order to prove the drug’s efficacy. In a heterogeneous treatment effect (HTE) setting where the drug has a clear positive effect on only a proportion of patients i.e. responders, traditional tests might be underpowered to detect the efficacy by testing for the difference in mean between the two groups. Our test could greatly benefit clinical trials because of the gain in power for detecting the drug’s heterogeneous effect.

## Supporting Information

Supplementary Figures and Tables

## Materials and Methods

### Statistical test for aberration enrichment

To calculate the enrichment score for a given variable of interest, we first sort the samples by that variable, then we walk through the list taking positive steps when we encounter a case and negative steps when we encounter controls. The enrichment score at a given position in the ranked list of samples, is a weighted sum of increments (for cases) and decrements (for controls): The formula for the *n^th^* position of the enrichment score *S_n_* is:

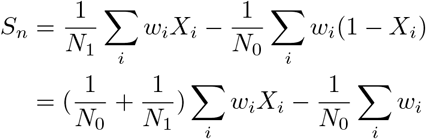

where *N*_1_ and *N*_0_ are the total numbers of cases and controls, **w** is the vector of weights, and *X* is the indicator variable describing which individuals are cases (1) and which are controls (0). The weights *w* can be the absolute values of the standardized levels of the variable of interest, thereby heavily weighting aberrations of large magnitude by making them correspond to larger steps in the walk. In practice we set the minimum on *w* to be 0.5 so that even individuals that are not outliers (near 0 standardized expression) can still contribute to our test. We chose to use this weighting scheme across all our experiments in order to make our test sensitive to the magnitude of aberration while still considering the ordering across all individuals when looking for an enrichment in cases versus controls. If we take *w* = 1 a constant across individuals, then only the ordering matters (similarly to Wilcoxon) and the scale no longer have an impact on the test statistic. This change of weighting scheme has a mild effect on the associations discovered in practice.

Under the null hypothesis, *X* is a random permutation of the case/control labels. The total number of cases and controls being fixed, the expectation and variance for the enrichment score at position *n* are:

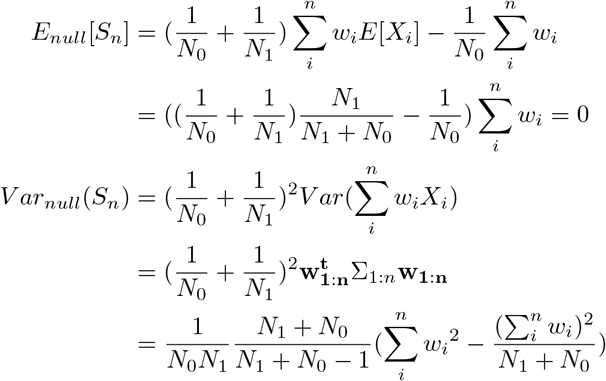

where Σ_1:*n*_ is the covariance matrix of *X*_1:*n*_

We standardize the enrichment scores by dividing them with the square root of the variance under the null *Var*(*S_n_*). All positions being comparable after standardization, we select the max standardized score across all positions as our test statistic 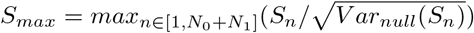.

The whole procedure described above is performed in both directions: sorting patients by increasing or decreasing expression levels and computing the max standardized enrichment score twice. Only the higher max standardized enrichment score is taken. in the example of gene expression data, this allows us to properly test for enrichment in overexpression and underexpression simultaneously. In a setting where only one direction of effect ought to be tested, we could perform the procedure on only one direction for a small gain in power.

We assess significance by running permutations on the case/control labels and observing how often we obtain a higher value for *S_max_*.

If we have an association, we can look back at which *S_i_* corresponds to *S_max_*. This will define the interval of values considered to be associated with cases versus controls. From there we can compute several useful interpretable quantities: such as the number of cases and controls in that interval, the odds ratio of being in that interval for cases, and the estimated *r* which is the proportion of cases affected (cases in that interval divided by all cases). We can also identify which individuals are affected and further study that group or enrich for it in future experiments. For example, in a clinical trial setting, our test would not only detect the heterogeneous drug effect but also return the identity of the patients who were affected (responders) which can be further analyzed and characterized.

### Real data preprocessing

The data downloaded from GEO is already in Log scale, we standardize it, apply PEER [63] to remove known confounders (if provided) and 30 hidden factors. Then we test the residuals for differential expression analysis. Genes with missing values and patients missing key clinical variables were removed from the analysis.

Our focus here is to find genes that follow the pattern of aberrant differential expression. Therefore we want to remove any broad signal in the data caused by confounders or hidden variables or the consequences of the disease itself (compensatory pathways, drug effects, etc). PEER helps us remove such broad signals. The same approach was used in previous work in order to detect expression outliers associated with rare eQTLs [33]. Note that PEER can potentially reduce the number of differentially expressed genes if some of the inferred PEER factors corresponds to broad effects of the disease status. Given that the hidden PEER factor can model pathways and transcription factors, it is also possible that some truly differentially or aberrantly expressed genes signal will be removed by PEER if these genes are regulating a large number of other genes’ expression. However, attempting the same analysis without removing any hidden factors resulted in a too large number of associations: in some datasets, almost all genes were significantly associated by any method (Limma, Wilcoxon, our test). This would make futile any attempt at recovering relevant mechanisms of disease. Therefore, we decided to go with the PEER hidden factor removal for the datasets analysed in this paper.

The Alzheimer data (GSE63063 [62]) contains the gene expression of 284 AD patients, 189 MCI (mild cognitive impairment) and 238 healthy matched controls measured in blood. It comes in two batches (USA and UK) using the Illumina HumanHT-12 V3.0 expression beadchip and Illumina HumanHT-12 V4.0 expression beadchip platforms respectively. We correct for gender, age, ethnicity, batch/platform in addition to the 30 PEER hidden factors.

The Parkinson data (GSE99039) contains the whole blood gene expression of 205 IPD cases and 233 controls, we corrected for gender and batch in addition to the 30 PEER hidden factors.

The inflammatory bowel disease data (GSE73094), contains the colon and terminal ileum gene expression of 608 samples from CD patients, 331 from UC patients and 50 samples from non IBD individuals. Overall 374 samples were taken during inflammation and 609 taken from non inflamed tissues. When we looked for association with CD versus UC and vice-versa, we used the non-inflamed samples and we selected individuals with one non-inflamed sample (a few individuals had multiple samples taken). This resulted in 181 UC samples and 314 CD samples. We corrected for group (code IBD2, IBD3, IBD4), tissue of origin, and 30 PEER hidden factors. When we looked for association to inflammation status, we took all samples (374 inflamed and 609 non inflamed) and we corrected for group, disease type and tissue in addition to the 30 PEER hidden factors.

The heritable breast cancer data (GSE47862) measures the gene expression in peripheral blood in 158 women with breast cancer and 163 controls. 226 women have a family history of breast cancer, 93 of which carry BRCA mutations. We corrected for cohort (Ontario or Utah) and 30 PEER hidden factors.

The breast cancer metastasis data (GSE48091), measures the gene expression in primary breast cancer tissue in 166 cases where metastasis happened and 340 cases without metastasis. We corrected for training/validation status and 30 PEER hidden factors.

The first schizophrenia methylation data (GSE74193) describes the DNA methylation in dorsolateral prefrontal cortex for 191 cases and 335 controls (after QC, removing duplicates and removing 8 cases whose reported gender differed from the predicted gender). We corrected for gender, race, batch, tissue composition in addition to the 100 PEER hidden factors.

The schizophrenia methylation data (GSE80417) measures the whole blood DNA methylation for 305 cases and 333 controls. We corrected for gender and age in addition to the 100 PEER hidden factors.

The Rheumatoid Arthritis methylation data (GSE42861) measures DNA methylation in peripheral blood leukocytes for 354 RA patients and 335 controls. We corrected for gender, age and smoking status in addition to the 100 PEER hidden factors.

The breast cancer miRNA data (GSE73002) describes the serum miRNA levels of 1280 breast cancer cases and 2686 controls. The ovarian cancer miRNA data (GSE106817) describes the serum miRNA levels of 399 ovarian cancer cases (including 79 borderline ovarian tumor) and 3647 non-ovarian cancer controls. The controls included 2759 healthy individuals and 859 individuals with other solid cancers (not ovarian). There were no additional clinical variables to use as confounders (In the ovarian cancer dataset, age had a missing value in the controls). In the classification experiment, we selected only the samples labeled ovarian cancer (320) and healthy controls (2759) for the ovarian cancer data. We corrected for 100 PEER hidden factors.

### Permutations analysis

To assess the significance of a considered variable’s max enrichment score, we run permutations on the case/control labels and we calculate how often the max standardized enrichment score generated by a permutation exceeds the score obtained on the real example. In order to compute accurate p-values, we need to run a very large number of permutations which can be computationally prohibitive. To reduce the number of unnecessary permutations we tried two approaches and found they lead to the same result.

Approach 1: For variables(e.g. genes) that are clearly not going to be significant, we do not need a high level of accuracy in estimating the p-value. Therefore, we adopt a gradual approach of successively running 100,1000,10000,100000,1000000,10000000,100000000 permutations only moving forward to the next step if less than 10 trials yielded higher test statistics than the real test statistic. We used this approach for all real gene expression datasets. By repeating the full experiment with more permutations every time, we essentially end up wasting a fraction of computations (the permutations from each previous cycle) but that is negligible compared to the gains in the majority of the genes where high accuracy is not needed to determine the clear absence of association. The factor 10 and the condition less than 10 trials work really well in practice, as we observed no bias in p-values estimations compared to the full permutation p-values.

Approach 2: When there is a large number of highly associated variables in a dataset, computing permutations for each variable can still be very slow using the first approach. By plotting, the test statistic and the log of the permutation p-values for all considered variables on the same graph, We observed that the function mapping test statistic and p-value is clearly monotonous and that the p-values could easily be predicted from the test statistic alone independently of the gene considered (See Supplementary Figure 5). This can be explained by the fact that the enrichment scores were standardized by accounting for each position and set of weights *w*, generating test statistics values (max over standardized enrichment scores) that are therefore comparable quantities across the genes/factors tested in the same dataset.

Under this assumption, there is no need to compute permutations separately for every gene/factor. Instead, we can keep the case/control labels and compute permutations on only one variable or even on a vector sampled from a Gaussian and then use those permutations’ test statistics to compute all the p-values from the test statistics (proportion of permutation test statistics that are higher than the true test statistic). We easily verify the quality of this approach by plotting the permutation p-values computed on only one Gaussian vector versus the p-values computed by doing permutations separately on every gene in Supplementary Figure 5. The correlation between the log p-values was 0.988. For datasets such as methylation data where there is a very large number of features, it is computationally prohibitive to compute permutations on every feature, especially given that a large number of permutations (10^8^) is needed for every feature to reach a useful level of accuracy for p-values (multiple hypothesis burden). The second approach was particularly useful in these datasets where approach 1 or full permutations were prohibitively slow.

Note that we do not assume that the mapping from test statistic to p-values is universal across datasets. Our second approach is applied separately on every dataset. The sample sizes and proportions of cases and controls are constant characteristics across phenotype permutations and could influence the mapping from the max standardized enrichment scores to p-values.

In our real data experiments, for the very significant genes/methylation sites/miRNAs with computed permutation p-value equal to 0 after 10000000 permutations (100000000 for methylation data), we estimated the plotted p-value by fitting a linear regression of log(p-value) as a function of a third order polynomial of the test statistic. We are interested in modeling the relation between the higher test statistics and the corresponding p-values. Therefore the regression model was fitted on the random permutations that reached a p-value of 0.05 or under among the set of all permutations performed. We verified that this procedure can lead to an accurate estimation of the low p-values, but in this paper we only used it for visualization purposes when plotting the 0 p-values variables in Figure 7-A. The significance of all novel associations reported was re-verified and was not affected by this approximation procedure since we only apply it on the very significant associations (*p* < 10^−7^, or *p* < 10^−8^ for methylation data).

### Classification of cancer versus healthy from miRNA data

We held-out an equal number of cases and controls from the data corresponding to 15% of the number of cases. We performed a cross validation on the remaining individuals. For each fold, we use our statistical test on the training data to perform feature selections then feed the selected and transformed features to an L1-logistic regression classifier (glmnet R package). We used the area until the Precision-Recall curve (AUPRC) to evaluate the performance. The hyperparameters of the classifier (here the regularisation λ) are chosen on the validation set and we report the performance of the model on the held-out test data (never seen by our test and by the classifier).

For feature selection, we use the same number of features for Limma and our method (300). For our method, we select only the heterogeneous features (aberration enriched) with *r* < 0.3. The optional feature transformation is based on the results of our test. For every feature, we determine the interval of aberrant expression driving the association. When the association is due to underexpression, this corresponds to all levels of expression lower than the expression level of the individual for which the max standardized enrichment score was reached. When the association is due to overexpression, this corresponds to all levels of expression higher than the level of the individual for which the max standardized enrichment score was reached (See Figure 1 for an illustration of this interval of interest). Once the interval of interest is defined, the feature transformation consists in assigning a value of zero to any individual outside that interval.

