## Supplement for "Finding associations in a heterogeneous setting: Statistical test for aberration enrichment"

### This PDF file includes:

Figs. S1 to S5
Tables S1 to S7

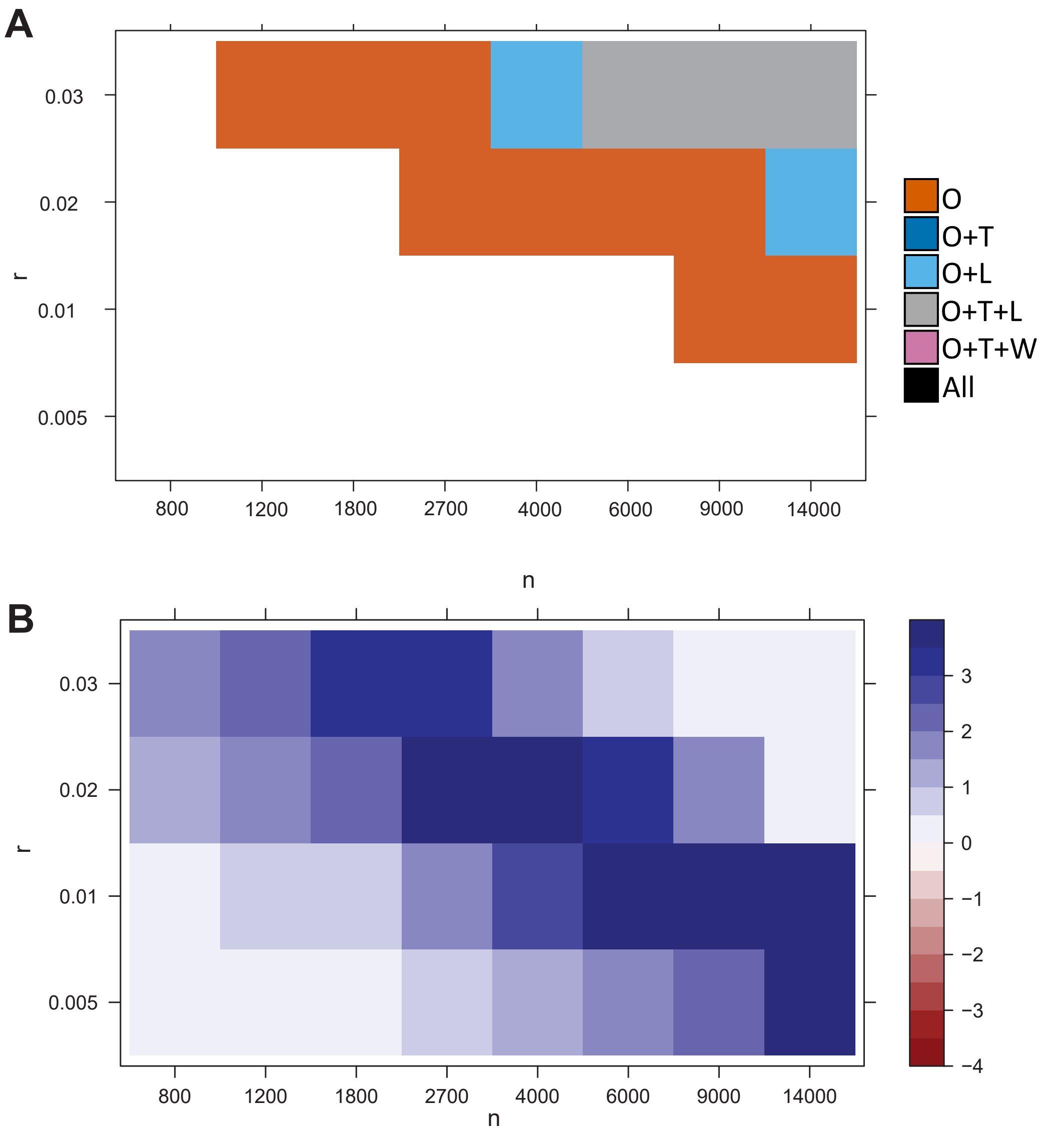


**Fig. S1.** **A** Ability of the different tests to detect the association for very low values of $r$. A method is able to detect the signal if the p-value is lower than the threshold in the the majority of 200 reruns. Here we show the results for $d = 3$. **B** Comparison of the p-value magnitude between our aberration enrichment test and the best out of t-test, Levene and Wilcoxon test. depending on simulations parameters $n$ (sample size) and $r$ (proportion of affected cases). Here we show the results for $d = 3$. The colors indicate the difference in log10 between the p-values returned. For example $2$ indicates that our test's p-value is two orders of magnitude (100 times) smaller than that of the best other method. We capped the maximal difference at $4$ for visual clarity. In reality the average difference in magnitude between our p-values and those of other methods can be much larger than 4 order of magnitude (10000 times).

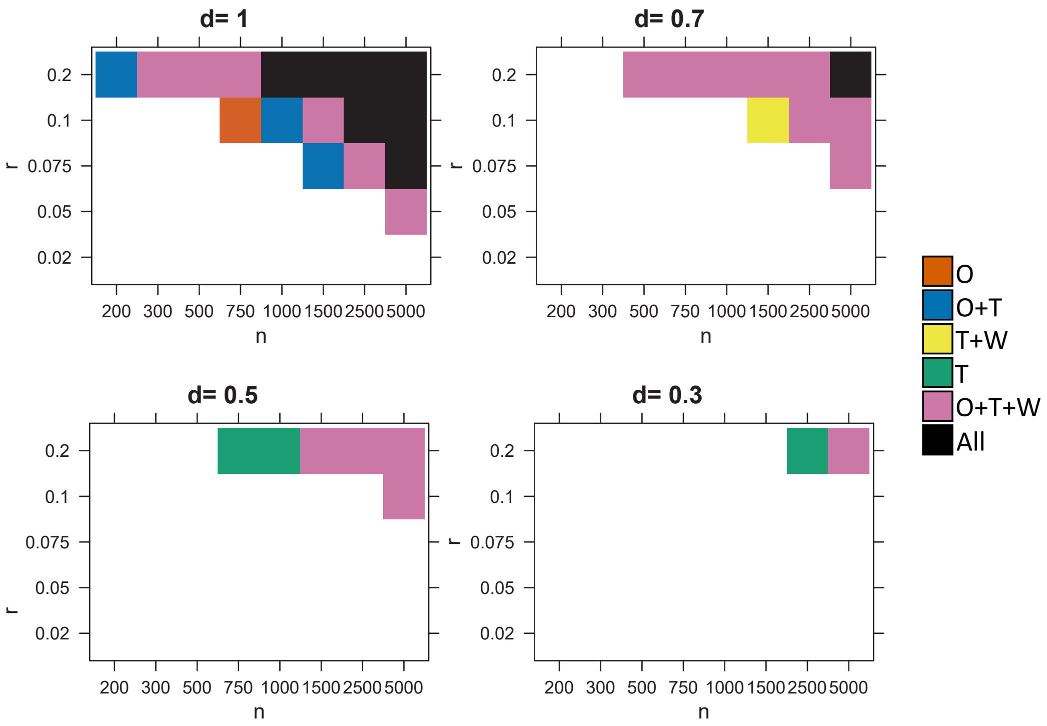


**Fig. S2.** Ability of the different tests to detect the association for low perturbation magnitudes $d$, depending on simulations parameters $n$ (sample size) and $r$ (proportion of affected cases). A method is able to detect the signal if the p-value is lower than the threshold in the the majority of 200 reruns.

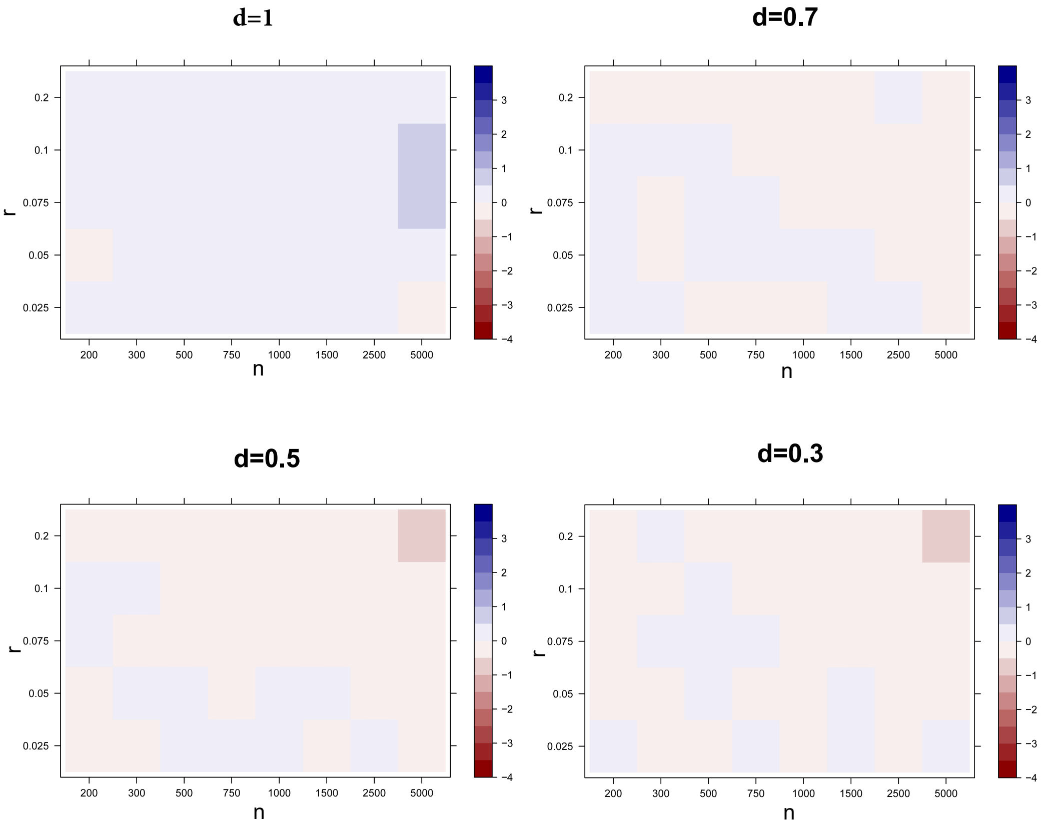


**Fig. S3.** Comparison of the p-value magnitude between our aberration enrichment test and the best out of all other methods for smaller sizes of perturbation $d$, depending on simulations parameters $n$ (sample size) and $r$ (proportion of affected cases). The colors indicate the average difference in log10 between the p-values returned by both method. 200 reruns were performed. Blue is for when our method is better than the best of the other methods (t-test, Wilcoxon and Levene)

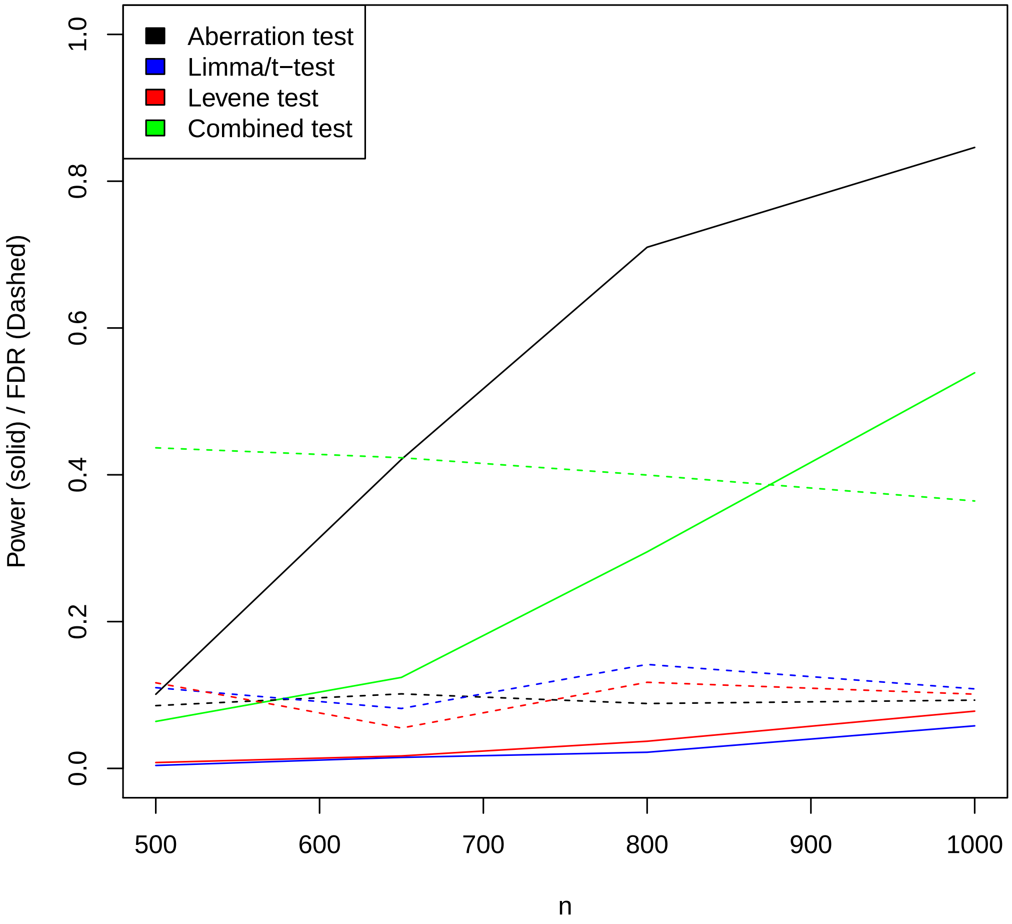


**Fig. S4.** False discovery rates (hashed) and power (solid) in the harder setting where $r = 0.05$ (proportion of affected cases) with larger sample sizes $n$. Ability of the different tests to detect the 10 simulated true genes among 25000. We fixed $d = 3$. We used an FDR threshold of $0.1$ for all methods. The average performance over 100 simulations is shown here.

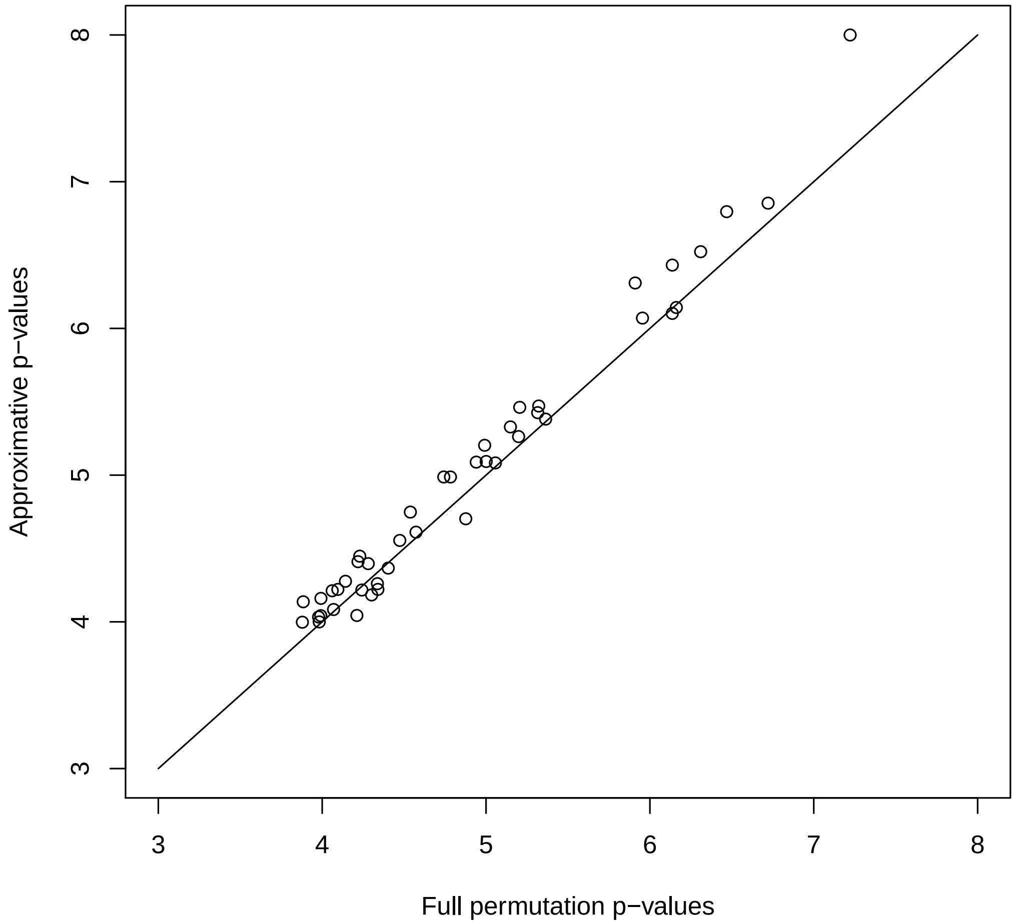


**Fig. S5.** Comparing p-values computed on a gene by gene basis with 100000000 permutations to p-values estimated with our optional approach of non-gene-specific p-value calibration (described in the method section as Approach 2). Both p-values computations were done on the same 50 probes from the Rheumatoid Arthritis Methylation data. The probes were randomly sampled from the top 500 most significant probes. Runtime was about a week on a personal laptop. 6 probes had p-values of zero by both estimation methods. For the remaining probes the correlation between log-pvalues was 0.988. The result indicates that the probes found significantly associated using the approximation were also significant by the full permutations and vice versa.

**Table S1. GEO Datasets used**

| ID | disease | Abv. | tissue | Data type | cases | controls |
|---|---|---|---|---|---|---|
| GSE63063 | Alzheimer disease | AD | blood | Gene expr. | 284 | 238 |
| GSE99039 | Parkinson | IPD | whole blood | Gene expr. | 205 | 233 |
| GSE73094 | Crohn's disease | CD | colon/terminal ileum | Gene expr. | 314 | 181 |
| GSE73094 | Ulcerative Colitis | UC | colon/terminal ileum | Gene expr. | 181 | 314 |
| GSE73094 | Inflammatory Bowel disease | IBD | colon/terminal ileum | Gene expr. | 374 | 609 |
| GSE47862 | hereditary breast cancer | - | peripheral blood | Gene expr. | 158 | 163 |
| GSE48091 | breast cancer metastasis | - | primary cancer | Gene expr. | 166 | 340 |
| GSE42861 | Rheumatoid Arthritis | RA | Leukocytes | Methylation | 354 | 335 |
| GSE74193 | Schizophrenia | Sch | prefrontal cortex | Methylation | 191 | 335 |
| GSE80417 | Schizophrenia | Sch | whole blood | Methylation | 305 | 333 |
| GSE73002 | Breast cancer | - | serum | miRNA | 1280 | 2686 |
| GSE106817 | Ovarian cancer | - | serum | miRNA | 320 | 2759 |

**Table S2. Effect of changing $k$ in the miRNA cancer datasets**

| Threshold | | Bonferroni | | | FDR < 0.2 | | |
|---|---|---|---|---|---|---|---|
| Method | k | our test | Limma | ∩ | our test | Limma | ∩ |
| breast miRNA | 30 | 480 | 0 | 0 | 658 | 1 | 1 |
| breast miRNA | 100 | 427 | 0 | 0 | 590 | 0 | 0 |
| breast miRNA | 200 | 425 | 0 | 0 | 580 | 0 | 0 |
| breast miRNA | 400 | 422 | 0 | 0 | 580 | 0 | 0 |
| ovarian miRNA | 30 | 462 | 58 | 49 | 1106 | 258 | 231 |
| ovarian miRNA | 100 | 364 | 5 | 4 | 984 | 31 | 31 |
| ovarian miRNA | 200 | 352 | 3 | 2 | 987 | 7 | 7 |
| ovarian miRNA | 400 | 361 | 2 | 2 | 975 | 6 | 6 |

**Table S3. Other omics: Number of genes detected with k=30**

| Threshold | Bonferroni | | | FDR < 0.1 | | |
|---|---|---|---|---|---|---|
| Method | our test | Limma | ∩ | our test | Limma | ∩ |
| RA methylation | 169 | 3 | 0 | 726 | 4 | 4 |
| Schizo. cortex | 67 | 0 | 0 | 326 | 0 | 0 |
| Schizo. blood | 266 | 0 | 0 | 28472 | 0 | 0 |
| breast miRNA | 483 | 0 | 0 | 628 | 0 | 0 |
| ovarian miRNA | 462 | 58 | 48 | 1012 | 197 | 179 |

**Table S4. Comparison with Wilcoxon: Number of genes detected. k=100**

| Threshold | Bonferroni | | | FDR < 0.1 | | |
|---|---|---|---|---|---|---|
| Method | our test | Wilcoxon | ∩ | our test | Wilcoxon | ∩ |
| RA methylation | 119 | 98 | 40 | 506 | 352 | 147 |
| Schizo. cortex | 22 | 18 | 6 | 139 | 85 | 21 |
| Schizo. blood | 75 | 61 | 7 | 530 | 242 | 58 |
| breast miRNA | 427 | 172 | 75 | 564 | 448 | 238 |
| ovarian miRNA | 364 | 34 | 22 | 849 | 310 | 193 |

Table S5. Gene expression data: Number of genes detected with k=100

| Threshold | Bonferroni | | | FDR < 0.1 | | |
|---|---|---|---|---|---|---|
| Method | esa | Limma | ∩ | esa | Limma | ∩ |
| Alzheimer vs ctr | 23 | 25 | 19 | 65 | 104 | 48 |
| Parkinson | 0 | 0 | 0 | 0 | 0 | 0 |
| CD | 2 | 0 | 0 | 2 | 0 | 0 |
| UC | 0 | 0 | 0 | 0 | 0 | 0 |
| IBD inflammation | 2 | 0 | 0 | 19 | 0 | 0 |
| breast cancer | 0 | 1 | 0 | 35 | 7 | 2 |
| breast cancer metastasis | 0 | 0 | 0 | 1 | 0 | 0 |

**Table S6. Comparison with Wilcoxon. Gene expression dataets. k=30**

| Threshold | Bonferroni | | | FDR < 0.1 | | |
|---|---|---|---|---|---|---|
| Method | our test | Wilcoxon | ∩ | our test | Wilcoxon | ∩ |
| Alzheimer vs ctr | 22 | 22 | 18 | 69 | 111 | 50 |
| Parkinson | 0 | 0 | 0 | 0 | 0 | 0 |
| CD | 2 | 1 | 1 | 2 | 3 | 1 |
| UC | 1 | 1 | 0 | 1 | 3 | 0 |
| IBD inflammation | 8 | 3 | 2 | 49 | 4 | 4 |
| breast cancer | 15 | 11 | 7 | 406 | 224 | 118 |
| breast cancer metastasis | 2 | 1 | 1 | 4 | 1 | 1 |

**Table S7. Reporting the false positive rate and the power of other tests. The average performance over 100 simulations is shown here.**

| n | 150 | 300 | 450 | 600 |
|---|---|---|---|---|
| Kolmogorov-Smirnov FDR | 0.07 | 0.09 | 0.055 | 0.13 |
| t-test FDR | 0.1 | 0.066 | 0.082 | 0.124 |
| Logistic regression FDR | 0.01 | 0.055 | 0.059 | 0.098 |
| ANOVA FDR | 0.01 | 0.055 | 0.059 | 0.098 |
| Levene test FDR | 0.09 | 0.102 | 0.08 | 0.11 |
| Wilcoxon FDR | 0.07 | 0.085 | 0.098 | 0.12 |
| Kolmogorov-Smirnov power | 0 | 0 | 0.003 | 0.01 |
| t-test power | 0 | 0.040 | 0.262 | 0.582 |
| Logistic regression power | 0 | 0.021 | 0.216 | 0.538 |
| ANOVA power | 0 | 0.021 | 0.216 | 0.538 |
| Levene test power | 0.002 | 0.070 | 0.358 | 0.739 |
| Wilcoxon power | 0 | 0.005 | 0.012 | 0.039 |
